# Bifurcation analysis of metabolic pathways: an illustration from yeast glycolysis

**DOI:** 10.1101/163600

**Authors:** Gosse Overal, Bas Teusink, Frank J. Bruggeman, Josephus Hulshof, Robert Planqué

## Abstract

In microorganisms such as bacteria or yeasts, metabolic rates are tightly coupled to growth rate, and therefore to fitness. Although the topology of central pathways are largely conserved across organisms, the enzyme kinetics and their parameters generally vary. This prevents us to understand and predict (changes in) metabolic dynamics. The analytical treatment of metabolic pathways is generally restricted to small models, containing maybe two to four equations. Since such small core models involve much coarse graining, their biological interpretation is often hampered. In this paper we aim to bridge the gap between analytical, more in-depth treatment of small core models and biologically more realistic and detailed models by developing new methods. We illustrate these methods for a model of glycolysis in *Saccharomyces cerevisiae* yeast, arguably the best characterised metabolic pathway in the literature. The model is more involved than in previous studies, and involves both ATP/ADP and NADH/NAD householding.

A detailed analysis of the steady state equations sheds new light on two recently studied biological phenomena in yeast glycolysis: whether it is to be expected that fructose-1,6-biphosphate (FBP) parameterises all steady states, and the occurrence of bistability between a regular steady state and imbalanced steady state in which glycolytic intermediates keep accumulating.

This work shows that the special structure of metabolic pathways does allow for more in-depth bifurcation analyses than is currently the norm. We especially emphasise which of the techniques developed here scale to larger pathways, and which do not.

## 1 Introduction

Metabolism is central to all life. The underlying network of enzyme-catalysed reactions changes with environmental conditions to sustain the living state. In microorganisms metabolic regulation is even more important, since metabolic rates are directly coupled to cellular growth rate, and hence to fitness. Understanding the dynamics of metabolic networks is therefore an important challenge in systems biology.

A comforting observation is that the reaction network of central metabolism has nearly identical topology across microorganisms. The enzyme kinetics and its parametrisation varies however between microorganisms, which prevents prediction of metabolic dynamics of poorly understood given the dynamics of those we understand much better. It is therefore of importance to develop methods that analyse the dynamics of unparameterised ordinary-differential-equation models of metabolic networks. That this is likely possible is because the mathematical functions that describe the rate of enzymes as function of the concentration of metabolic intermediates (reactant and effector molecules) and kinetic parameters are strongly constrained by protein biochemistry. Here we undertake this challenge for glycolysis, the most conserved metabolic pathway in biology. We focus on its implementation in the yeast *Saccharomyces cerevisiae*, for which we know its dynamic behaviour (such as oscillations, bistability and exploding states) and plenty of enzyme kinetic information is available.

The glycolysis pathway has been the focus of research for decades. It metabolises glucose into pyruvate, thereby using the free energy to generate 2-adenine 5’-triphosphate (ATP) and the freed electrons to reduce nicotinamide adenine di-nucleotide (NAD) to NADH. Glycolysis is essential for cells: it provides much of the ATP that drives countless biological processes, and glycolysis provides some of the most important precursor molecules, such as pyruvate, from which amino acids, lipids and other macromolecules are synthesised. Moreover, many branches feed into glycolysis, so that other sugars, such as fructose, galactose, sucrose, maltose, lactose and others, may be metabolised through this pathway as well.

When yeast is deprived of oxygen, its glycolysis converts pyruvate further into ethanol and CO_2_ by oxidising NADH. This yields a very fast but inefficient energy production, in which 2 out of the potential 12 ATP are obtained from one molecule of glucose. The yeast glycolytic pathway has been studied extensively over the years, and two fully detailed models have been developed which include fully parameterised reaction kinetics for all the individual enzymatic steps [20, 9]. Nevertheless, despite this wealth of detail, a number of regulatory and dynamical aspects of yeast glycolysis remain poorly understood at present.

When the metabolite concentrations external to the cell change, for instance if a new food source becomes abundant, the cell’s limited enzyme production needs to be redistributed by the gene regulatory network to reach a new steady state to maximise the flux through glycolysis. It has very recently been shown that enzyme levels are indeed pervasively tuned to maximise growth rate [10]. The gene network is responsible for tuning enzyme levels, but it needs input from the pathway it controls to sense changes in the environment. Nutrient-specific membrane receptors could provide such input, and yeast has a detailed glucose-sensing mechanism [4]. Nevertheless, as in most bacteria [11], yeast cells also sense the flux through glycolysis by using glycolytic intermediates binding to transcription factors. These then influence gene expression. Experimental evidence suggests that the glycolytic intermediate fructose-1,6-biphosphate (FBP) acts as a flux-sensor [2, 8], directly influencing the gene network and thereby inducing changes in the glycolytic enzyme levels, a form of adaptive control [15].

However, it less clear why FBP should play this role as sensor. For FBP to function properly, its concentration should contain sufficient information to assess the metabolic flux through glycolysis. The FBP concentration should therefore be associated to a unique steady state concentration profile. This has been shown to be true experimentally and a mechanism has been proposed [11], but it is much less clear how this property emerges from the kinetic properties of the glycolytic pathway. We investigate here for a detailed core model under what parameter assumptions FBP indeed parametrises steady states. We also ask the question whether the steady states may actually be faithfully predicted by a flux value, for instance one of the FBP-consuming fluxes.

Glycolysis, in yeast specifically, has held another mystery for years. Yeast can synthesise trehalose from the glycolytic intermediate glucose 6-phosphate. This reaction is not a step of the glycolysis pathway, so one does not expect glycolysis to fail when this reaction is disabled by means of a gene knockout. However, many cells of the mutant in which this particular knockout is performed, the tps1-Δ mutant, are not able to grow on glucose [22]. In [22] it was revealed that this mutant shows a form of bistability between a regular steady state and an imbalanced state in which some intermediate metabolites, including FBP, accumulate in the cell, reaching toxic levels. In fact, also wild type yeast suffers this problem, but only a small part of the wild type population enters the imbalanced state [22]. The trehalose branch does not completely inhibit this effect, but makes it less likely for glycolysis to fail and more likely to grow well; in dynamical systems terms: the basin of attraction of the imbalanced state is reduced in size, so that the regular steady state is reached from a wider range of initial conditions. In a small core model of yeast glycolysis with three variables [14], this bistability was studied in detail. Here we improve upon those results by studying a more detailed core model which contains five variables and seven reaction fluxes.

More generally, we aim to develop analytical techniques which exploit the special structure of metabolic pathway ODEs, and which may also give general insight into the bifurcation structure for larger pathways. Such “parameter-free analyses” are usually performed on core models of up to three variables or so. It is currently beyond our ability to prove nearly anything about fully detailed and well-characterised pathway models such as the ones by Teusink *et al.* and Hynne *et al.* [20, 9], and insight is generally only obtained by numerical simulations using measured parameter values. In most cases, however, the situation is worse: many, if not most, kinetic parameters are unknown, making it difficult to decide their value in simulations. We therefore highlight in the Discussion which of the analytical techniques developed in this paper scale to larger networks.

### Introducing the glycolytic pathway

Glycolysis is split into two lumped reactions (see Fig. 1): upper glycolysis (*v*_1_), using glucose and 2 ATP (*a*) to produce FBP; and lower glycolysis (*v*_2_), using FBP (*f*) to produce 2 pyruvate (*y*), 2 NADH (*n*) and 4 ATP. This modelling of upper and lower glycolysis is done to be able to focus on the imbalanced state, when the lower part cannot keep up with the upper part and FBP accumulates. The reaction *v*_4_ uses NADH and pyruvate to produce ethanol. This reaction, when combined with glycolysis, makes up ethanol fermentation.

**Figure 1:**
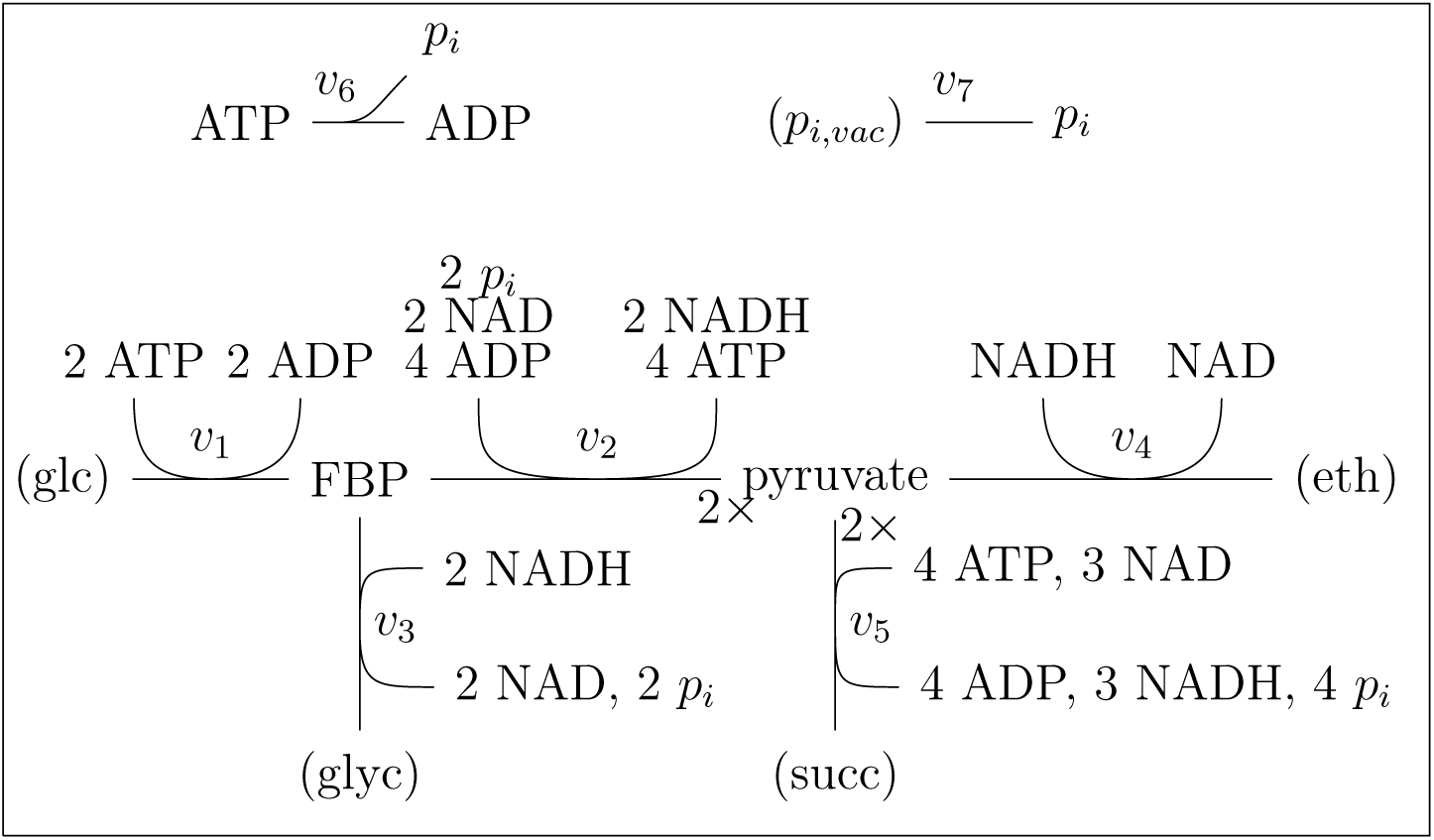
A graphical representation of the stoichiometry of our model. The nodes are the different metabolites, the arrows are the reactions. When a metabolite species is between brackets, the con-centration is disregarded or assumed constant in the model.

There are two side branches from the main pathway: one producing glycerol (*v*_3_) from FBP and 2 NADH, and one producing succinate and 3 NADH (*v*_5_) from 2 pyruvate and 4 ATP. They are included to allow redox balance and also carry a significant flux in experiments [20].

The experimental setup connected to the model is balanced growth of the population after starved yeast is presented with abundant glucose. The glucose will of course eventually be depleted. However, the restricted timeframe of the experiment is when yeast is growing exponentially. Therefore we can assume the concentration of glucose to be constant over the relevant time frame. The concentrations of ethanol, glycerol, and succinate are disregarded with the assumption of product insensitivity of *v*_4_, *v*_3_ and *v*_5_ respectively.

Aside from this pathway and its side branches, a general linear ATPase (*v*_6_) is included to model the total ATP use of the cell and balance the production by the glycolytic pathway. Inorganic phosphate, *p*_*i*_ (*p*), is a substrate and product of some of these reactions; for its stoichiometry, see Figure 1. Furthermore, the *p*_*i*_ concentration is dynamically buffered (*v*_7_), which corresponds to diffusion between the cytosol and the vacuole. We assume that the concentration inside the vacuole is not influenced on our time-scale and is constant (π). Therefore, *p* will be steered towards π by *v*_7_, the concentration of inorganic phosphate in the vacuole.

Conservation laws dictate the concentrations of ADP and NAD. The total concentration of ATP and ADP is constant (*a*_*T*_) and so the ADP concentration (*a*_*T*_ *- a*) is a dependent variable. Likewise the NAD concentration (*n*_*T*_*- n*) is a dependent variable. The parameters *a*_*T*_ and *n*_*T*_ are determined by the initial conditions.

The reactions in the model are lumped, and therefore we cannot use the detailed rate functions given in the Teusink or Hynne models [20, 9]. Instead, we have chosen Michaelis-Menten type dynamics (see Figure 3). These have the property to be monotone increasing in the substrate concentrations, a property that will be greatly exploited in our analysis. The reaction *v*_1_ in our model corresponds to phosphofructokinase (PFK), a complex enzyme with many binding sites for allosteric activation and inhibition. We simplify here by assuming that *v*_1_ depends only on *a* [6]. Despite this simplification, PFK still catalyses the most complex reaction: as a function of *a*, PFK is not always monotone. The reaction flux increases for small *a*, because ATP is a substrate, but at some point decreases, because ATP also allosterically inhibits PFK (see Figure 2 for a sketch).

**Figure 2:**
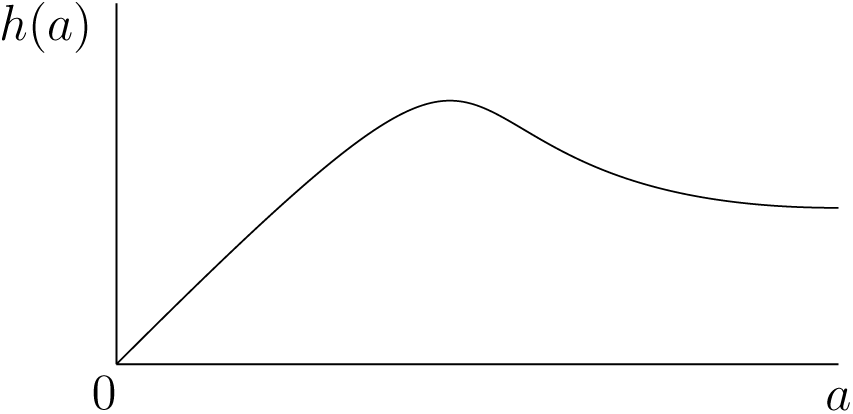
Schematic illustration of *h*(*a*), with *h*^*'*^(0) = 1 i.e. *v*1*'* (0) = *V*_1_.

**Figure 3:**
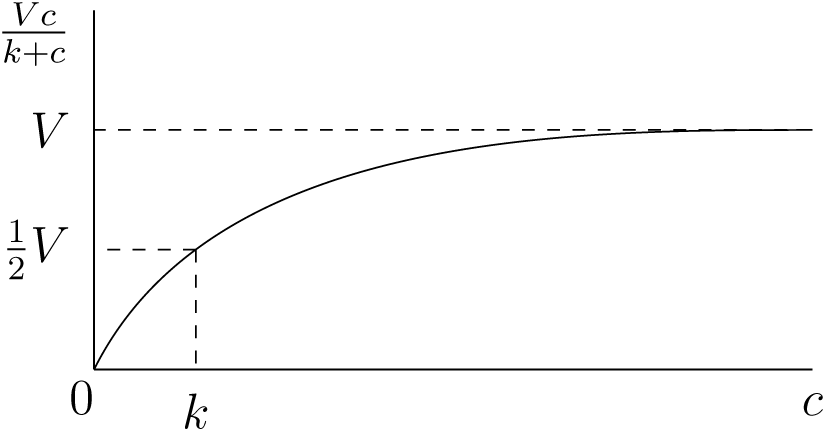
Michaels-Menten kinetics for substrate concentration *c* with parameters *V* and *k*.

### A first overview of the main techniques and results

In this section we only introduce the structure of the model, delaying a full description to Section 2, and give a first overview of the main techniques and results.

The independent variables of the model are the concentrations of FBP (*f*), *p*_*i*_ (*p*), pyruvate (*y*), ATP (*a*) and NADH (*n*), which are collected in the vector ***x***, and 7 reactions *v*_1_, *…, v*_7_ collected in ***v***(***x***).

The model is a system of differential equations,

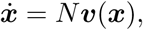

with stoichiometric matrix *N* and reaction rates ***v***(***x***) detailed in Section 2.

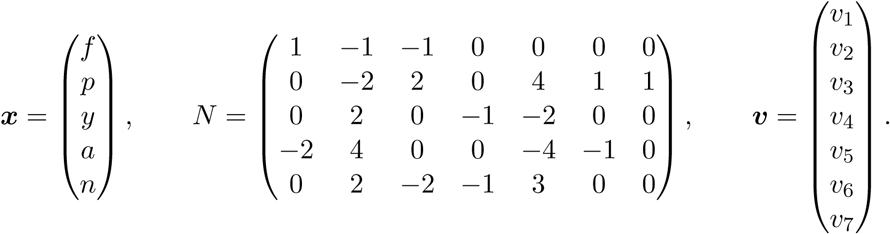

Each row of *N* denotes how many molecules of that metabolite are used as a substrate (negative entries), or produced (positive entries), by the 7 reactions (compare with Figure 1).

As explained before, we are particularly interested in the steady states. We will classify the families of steady states in terms of a natural bifurcation parameter.

To make it easier to solve the steady state equations *N* ***v***(***x***) = **0**, we would like the different reactions to be isolated in these equations, i.e. have many reactions appear in only one equation and each equation have as few reactions as possible. Mathematically, any basis of the row space of *N*, Row *N*, yields the same null space and therefore the same steady states. In Section 3.1 we will give a general method to find an appropriate basis which separates the reactions in the steady state equations. One example set of equations coming out of this procedure here is

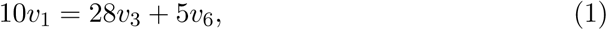

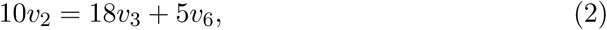

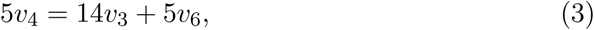

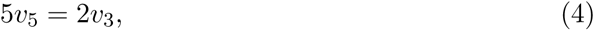

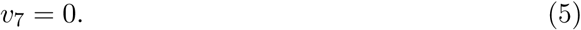

Note that *v*_1_, *v*_2_, *v*_4_, *v*_5_ and *v*_7_ each occur in only one equation. This system is equivalent to *N* ***v***(***x***) = **0**.

The bifurcation parameter should correspond to a biological quantity that can be changed in experiments. For the experimental setup, a food source such as glucose has to be provided, meaning that the concentration of the food source can be varied as a parameter. Looking at our model the influence of nutrient levels corresponds to the *V*_*max*_ of *v*_1_. Choosing this as our bifurcation parameter *λ*, it solves the steady state equation (1),

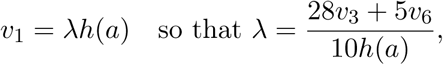

with *h*(*a*) a non-monotone function of *a* detailed in Section 2. So given a solution to equations (2)–(5), in terms of ***x***, equation (1) yields a unique *λ* and therefore gives us a direct description of the bifurcation curve. In this way the challenge PFK (*v*_1_) poses to analysis (its non-monotonic behaviour in *a*) is tackled. Moreover, even if *v*_1_ would have a more complicated form, e.g. if *h* would depend on multiple variables, *λ* would still solve equation (1) as above.

The equilibrium states, where there is zero flux, are shown to be two axes of the phase space and at the intersection of these there is a (complicated) bifurcation of a simple eigenvalue. We provide an explicit expansion of the emergent curve of steady states into the extended phase space 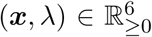 (Section 3.2). We prove that all non-trivial solutions to the steady state equations are locally described by a single, one-dimensional curve and that the FBP concentration parameterises this curve under a mild parameter condition (Section 3.3). This result follows from the Implicit Function Theorem considered for the steady state equations (2)–(4). This means for instance that pitchfork bifurcations are excluded.

The steady state equations (2)–(4), are all nonlinear in the variables ***x***, but they are of course linear in ***v***. A suitable coordinate transform in which the ***x*** variables are replaced by ***v*** variables makes most of this linearity, and should facilitate analysis [14]. Such a coordinate transform may be explicitly calculated in our case, together with its inverse transform (Section 3.4). The transformation heavily relies on the monotonic dependence of the reactions *v*_2_, *v*_3_, *v*_4_ *v*_6_ and *v*_7_ on their respective variables. The strategy may be summarised as follows. Equations (2) and (3) allow *v*_2_ and *v*_4_ to be expressed as linear combinations of *v*_3_ and *v*_6_. Apart from the trivial (5), this leaves (4). We express *v*_5_ in (4) as a function of *v*_2_, *v*_3_, *v*_4_ and *v*_6_ using the coordinate transform, and then replace *v*_2_ and *v*_4_ by their respective linear combinations of *v*_3_ and *v*_6_. The remaining problem is hence reduced to solving only one equation, (3), in two variables, *v*_3_ and *v*_6_. The resulting solution is substituted into (1) to produce the bifurcation curve.

Solving the last steady state equation in *v*_3_ and *v*_6_ involves some tedious work which we manage to overcome, to conclude that for a wide range of parameters the steady states are indeed parameterised by *v*_3_, corresponding analogous to parameterisation by the FBP concentration (Section 3.5).

This method of rewriting the steady state equations into nonlinear equations for ***v*** also allows us to study imbalanced states, in which for instance *f → ∞*: then *v*_3_ is still finite, and the imbalanced states form regular steady states for the flux variable system. (This was in fact the original motivation to introduce such coordinate transformations [14].)

To summarise, the steady states form a single branch in the extended space with *λ* and the five variables. The branch starts in a specific equilibrium state and, for a wide range of parameters, connects to an imbalanced state with infinite FBP. From this point at infinity, we may study the “system at infinity” to uncover other imbalanced states at different values of *λ*, and hence bistability between regular steady states and imbalanced ones (Section 3.6).

## 2 The mathematical model

In complete detail, the system of equations is

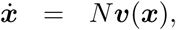

where ***x***, *N*, and ***v*** are

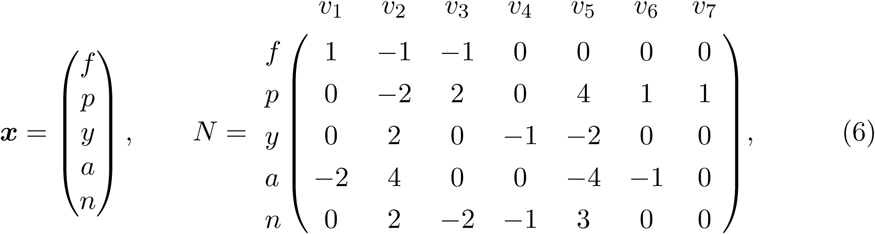

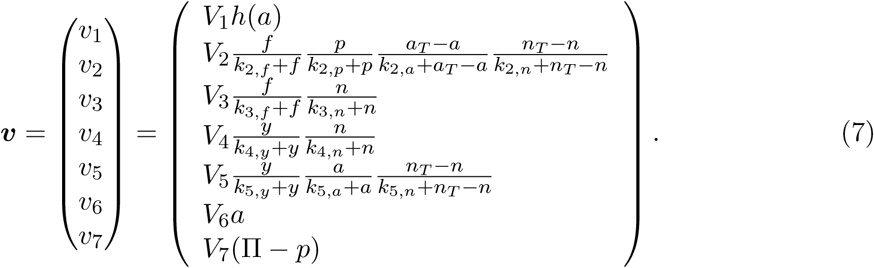

For the variables in ***x***, we demand

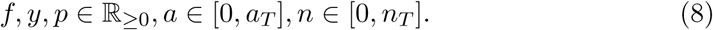

In the definitions of the reaction rates, all parameters are positive and *h*(*a*) is defined as follows (see also Figure 2 for a sketch),

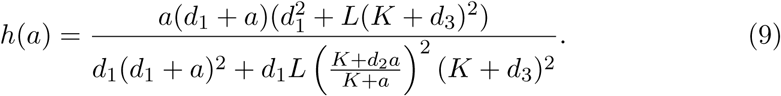

The parameters *d*_1_, *d*_2_, *d*_3_, and *d*_4_ are also positive. This formula is based on [20], where the concentrations of all metabolites apart from *a* are assumed to be constant [6], and *V*_1_ is rescaled such that *h*^*'*^(0) = 1.

The bifurcation parameter is *λ* = *V*_1_ *≥* 0.

## 3 Steady state analysis

The goal of this section is to gain insight into the steady states; steady states are solutions ***x*** to the steady state equations, given by

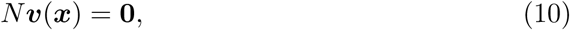

with ***x***, *N*, and ***v*** defined as in (6) and (7). We aim to solve these equations for ***x*** and the bifurcation parameter *λ* = *V*_1_.

## 3.1 Suitable representations of the null space of *N*

The solutions for ***x*** must satisfy (10). Thus ***v***(***x***) must be in the null space of *N*, Nul *N*, which is perpendicular to Row *N*. In order to get alternative steady state equations, we need a basis of Row *N*; the independent rows of *N* can serve as such a basis, but any independent set of five vectors in the row space will define the same null space. Choosing a new basis intelligently will generate steady state equations more amenable to analysis, with the same solution space as (10). We will provide a general method to construct such bases.

For the steady states in terms of the reactions ***v*** there is a well-established theory. The steady state flux distribution cone is the set of all possible solutions ***v*** to equation (10). This cone can be described by the Elementary Flux Modes (EFMs) [17, 18, 13]. This theory is largely based on linear algebra, because the cone is contained in the null space of *N*, but there are also positivity constraints: if a reaction is irreversible, it can only have positive flux.

We provide a brief summary of the theory on flux modes to keep this work self-contained. The interested reader can find a more detailed overview in [17, 18, 13]. A flux mode is represented by some fixed, nonzero reaction vector ***V***, which is a solution to (10) and another vector ***v*** is part of this flux mode if and only if there exists some positive constant δ ∊ ℝ_≥0_ such that ***v*** = *δ****V***. So a flux mode has fixed ratios between fluxes. An Elementary Flux Mode is, loosely speaking, a flux mode with a maximal number of zero entries. The steady state flux distribution cone is the convex set of all flux modes. There is a competing concept to describe the flux modes by means of extremal pathways [16], which also span the cone. The smallest set of flux modes such that every flux mode is a convex combination defines the set of extremal pathways. Every extremal pathway is an Elementary Flux Mode, but there are Elementary Flux Modes that are not extremal pathways. These appear if there is some reversible reaction that has a positive entry in one extremal pathway and a negative entry in another. The specific convex combination of those extremal pathways, where the reversible reaction has a zero entry, yields an Elementary Flux Mode, because it has an extra zero, but it is not an extremal pathway, because it is a convex combination of two extremal pathways.

When considering equation (10) as a problem for ***v***, the positivity constraints of irreversible reactions have to be imposed, yielding the inequalities that put EFM theory beyond linear algebra, into linear programming. However, if ***x*** is a vector of admissible concentrations, these constraints follow naturally, because the functions in ***v***(***x***) will satisfy them by definition (7). Therefore EFM theory is not necessarily the primary tool when the dynamics of ***x*** are considered, but it still leads to insight: if ***x*** is a steady state, ***v***(***x***) will be in the steady state flux distribution cone as a positive combination of the EFMs.

Besides this insight, the EFMs can be useful for the linear problem: the linear space that is spanned by the EFMs is exactly Nul *N*, so the space perpendicular to all EFMs *is* Row *N*. Every EFM has the property that the number of zero entries is maximised. This yields a constructive way of finding representative equations that define the steady state in which fluxes appear most sparsely. It uses a simple proposition from linear algebra.

**Proposition 1.** *If N is an n × m stoichiometric matrix and* ***m***_1_, *…,* ***m***_*k*_ *are the Elementary Flux Modes, then for a* ***w*** *∈* R*m,*

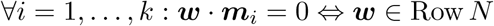

*Proof.* ***m***_1_, *…,* ***m***_*k*_ are such that they span the null space of *N*, therefore

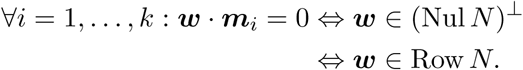

We apply this proposition to our model. Calculation of EFMs for particular pathways may be done using algorithms such as for instance *efmtool* [1] for MATLAB.

**Corollary 2.** *Let N be given by* (6). *Then* Row *N is the space perpendicular to the EFMs of our model, given by*

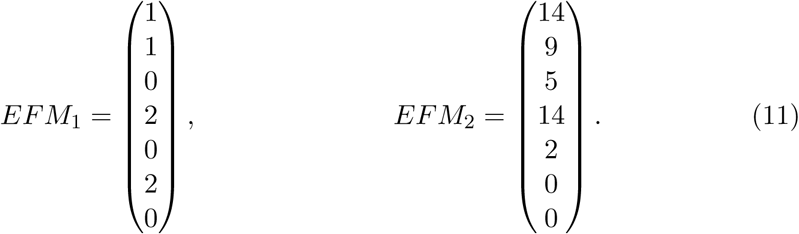

As noted in the introduction, the equations are most easily solved if some reactions only appear in a single equation. We will construct a matrix which has the same row space as *N* using Corollary 2.

We start by finding a diagonal submatrix in the matrix (*EFM*_1_, *EFM*_2_), e.g. finding those fluxes that are only represented in one EFM and have a zero entry in the other EFM. The only nonzero entry of *EFM*_1_ that has a zero entry in *EFM*_2_ is *v*_6_, while *EFM*_2_ has both *v*_3_ and *v*_5_ nonzero, but they are zero in *EFM*_1_. Thus there are two choices, *v*_3_ and *v*_6_, or *v*_5_ and *v*_6_.

Starting with a 5 *×* 5 identity matrix and inserting two columns in-between, such that they become the columns at the fluxes we have chosen, we construct a 5 *×* 7 matrix. The inserted columns are chosen such that every row is perpendicular to both EFMs in (11):

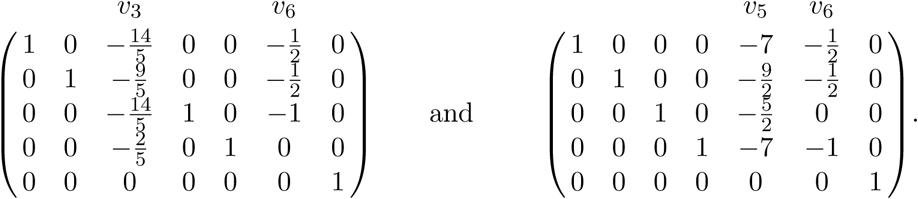

We are able to compute each entry in those two special columns individually because of the special choice of *v*_3_ and *v*_6_ (or *v*_5_ and *v*_6_). By multiplying the rows of the above matrices with the denominators of their entries, we get the following integer matrices,

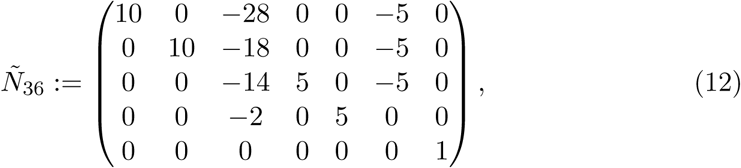

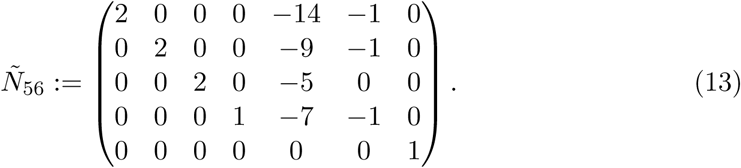

Understanding this construction, we see that we can be more creative. In fact, we may now also pick and choose rows from both these matrices, in such a way that the resulting new matrix still has rank 5. So for example, we could use *Ñ*36 to choose equations for *v*_2_ and *v*_4_ (and *v*_3_ and *v*_6_), but *Ñ*56 to complete the system with equations for *v*_1_ and *v*_3_ (and *v*_5_ and *v*_6_):

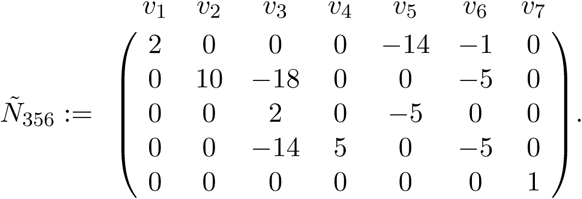

The difference is in the distribution of the reaction rates across these five equations. In *Ñ*36 and *Ñ*56, one reaction rate appears in four out of five equations, and four reaction rates only once. In *Ñ*356 reaction rates appear maximally three times, but only three reactions appear once.

The matrices *N, Ñ*36, *Ñ*56 and *Ñ*356 all have the same row space, since we constructed their rows to be independent and perpendicular to (11) (Corollary 2) and the matrices have equal rank. Thus the following problems are equivalent for ***v***(***x***),

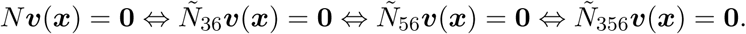

In this paper we use all three new matrices, making use of their individual properties to answer specific questions.

### 3.2 The equilibrium states

The system has trivial solutions, which we refer to in this work as equilibrium states.

**Definition 3.** *A state* ***x*** *is called an <u>equilibrium state</u> if* ***v***(***x***) = **0**.

These states, trivially satisfying *N* ***v***(***x***) = **0**, are steady states. In other models, chemical equilibria are usually dynamical steady states, where the forward and backward reactions are balanced, but in our model nearly all reactions are irreversible (the exception being *v*_7_). This modelling choice is based on the assumption that the glycolytic flux is much greater than the reverse, therefore the backward reactions can be disregarded. As a modelling artifact, the equilibrium states in our model are not balanced in forward and backward reactions, but have zero forward and zero backward reactions. As such, they cannot represent a living cell. Instead, the interpretation should be that, if in our model the equilibrium state is stable and the non-equilibrium state is unstable, the cell cannot use glycolysis for its ATP needs, but must turn to other pathways. On the other hand, if in our model the equilibrium state is unstable and there is a non-equilibrium steady state that is stable, then the cell can use glycolysis for its ATP needs. We will show that there is a specific value of our parameter *λ*, where we switch between these two situations in a transcritical bifurcation.

#### Lemma 4.

*The equilibrium states are 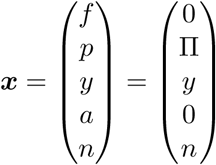, where yn* = 0

*Proof.* The reaction functions (7) have a product structure where many factors cannot be zero, including all denominators and parameters. Division by these nonzero factors yields

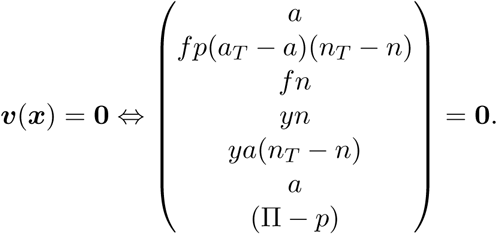

This is equivalent to

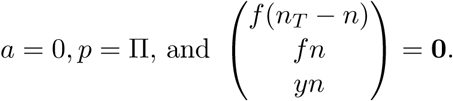

It follows that *fn* = 0 = *f* (*n*_*T*_ *- n*), therefore *f* = 0.

We conclude that

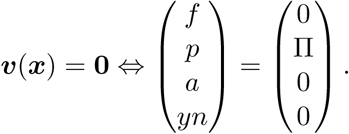

So *n* = 0 or *y* = 0 is sufficient. Thus the equilibrium states are a family of two lines on the boundary of the metabolite space. Either (0, π, 0, 0, *n*), where *n ∈* [0, *n*_*T*_] or (0, π, *y,* 0, 0), where *y ∈* ℝ_*≥*0_.

The intersection of these two families is the equilibrium

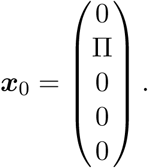

In the rest of this section we will show that ***x***_0_ is, in fact, the only relevant equilibrium, because a transcritical bifurcation occurs at ***x***_0_, giving rise to the non-equilibrium steady states. We also show that any non-equilibrium steady state that is near the equilibrium states is on this emergent curve. Furthermore, at this bifurcation ***x***_0_ transfers local stability to the regular non-equilibrium steady states.

The equilibrium state ***x***_0_ is thus a highly degenerate point, where three families of equilibrium states meet. The bifurcation analysis for this point is therefore quite subtle, and is treated in detail.

Note that all equilibrium states have *a* = 0; reactions *v*_5_ and *v*_6_ have a factor *a* in their definitions (7). A natural choice for the steady state equations is then

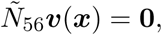

where Ñ_56_ is defined in (13), because it has the concentration *a* as a common factor in the negative parts of the equations.

We are interested in finding a transcritical bifurcation of the equilibrium state ***x***_0_ and proving that no other steady states exist close to the equilibrium states. Assume therefore that ***x*** is in a suitable neighbourhood around the equilibrium states, such that *a*_*T*_ *- a >* 0 and all denominators in the definition of ***v***(***x***) (7) are positive. Then we can introduce some simplifying notation. The steady state equations are

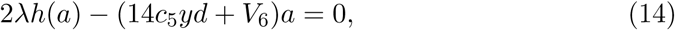

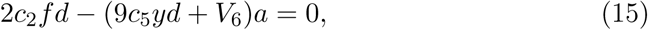

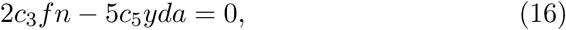

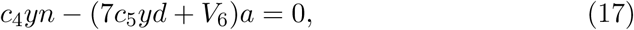

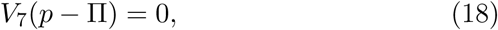

where *d* = *n*_*T*_ *- n* and

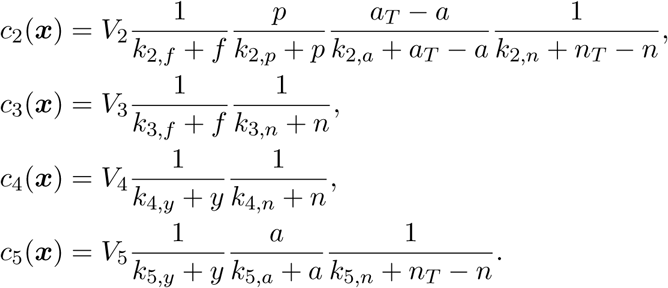

In other words, function *c*_*i*_(***x***) *>* 0 denotes all terms of reaction *v*_*i*_ that are nonzero in this neighbourhood. In this way, we can see more clearly the terms that yield equilibrium states and dividing out those terms, we find the equation for the non-equilibrium steady state curve. To keep the expressions transparent, we often drop the dependence of *c*_*i*_ on ***x***. Moroever, any steady state will have *p* = π, so we assume this for the rest of this section.

Equations (17) and (15) can be rewritten as

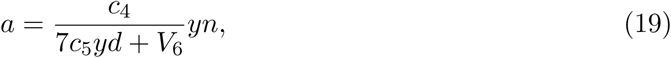

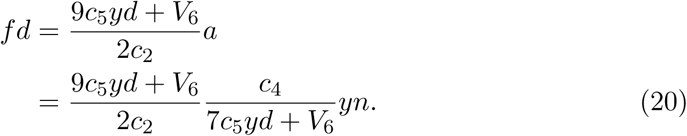

We multiply equation (16) with *d*, such that we can insert the solutions (19) and (20) for *a* and *fd* respectively,

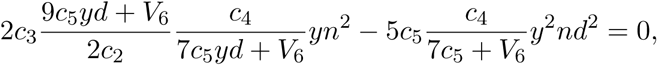

which simplifies to

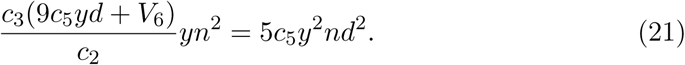

So it follows that *y* = 0, *n* = 0 or

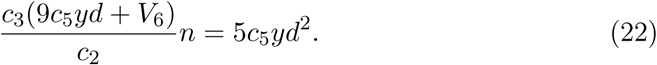

Together with equations (19) and (20), having *y* = 0 or *n* = 0 means that we have an equilibrium state as *f* = *a* = *yn* = 0.

The term *d* = *n*_*T*_ *- n* was not included as a factor in *c*_2_ or *c*_5_, since the equilibrium state

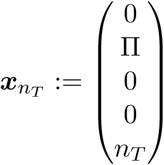

was still part of our neighbourhood of equilbria. In other words, *d* = 0 is still a possibility, but the analysis requires strict positivity of *c*_2_ and *c*_5_. Now we will show that around this specific equilibrium state ***x***_*n*_*T*, there are no other steady states than equilibrium states. In fact, we show that if (21) holds, then (22) cannot hold suffciently close to ***x***_*n*_*T* and that we can only have *y* = 0.

The equilibrium state ***x***_*n*_*T* has *y* = 0 and *d* = *n*_*T*_ *- n*_*T*_ = 0. If we now restrict to an even smaller neighbourhood, namely a ball around this equilibrium state, we see that both *y* and *d* have absolute values smaller than the radius of the ball. The concentration *n* however is close to *n*_*T*_, so the lhs of equation (22) is nonzero and does not get smaller as we decrease the size of the ball, while the rhs decreases with the third power of the radius of the ball. Hence if we take a small enough ball, equation (22) cannot hold and we conclude that around ***x***_*n*_*T*, if the steady state equation (21) holds, it yields *y* = 0. So the only steady states in this ball are equilibrium states.

For equation (22) to hold, *n* and *y* must be of the same magnitude, so let us assume for the rest of this argument that we are in a suitable neighbourhood of ***x***_0_, where *y* and *n* are small.

The term *h*(*a*) has concentration *a* as a factor, so we can write it as 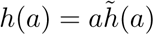, where 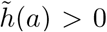 can be inferred from the definition of *v*_1_ (9). Moreover, *h*(*a*) was defined such that *h*^*'*^(0) = 1, and since 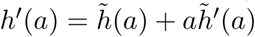,

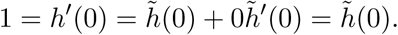

Equation (14) can be rewritten as

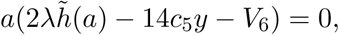

which gives us *a* = 0 or

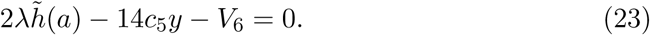

So there are two possibilities, either *yn* = *a* = *f* = 0, which means we have an equilibrium state and *λ* can have any value, or the equations (19), (20), (22) and (23) hold. The equations (19), (20), (22) and (23) are the (non-trivial) steady state equations. Equations (19), (20) are directly equivalent to the original steady state equations (15) and (17), but in (22) and (23) the degenerate equilibria *y* = *n* = 0 in (14) and (16) have been divided out. These four equations (19), (20), (22) and (23) can be rewritten as

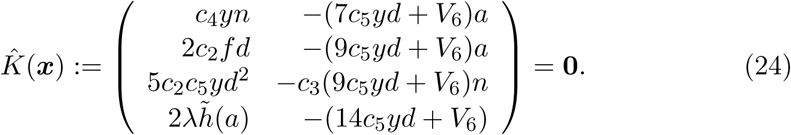

The equilibrium state ***x***_0_ is a solution for equation (24). Using 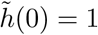, we have

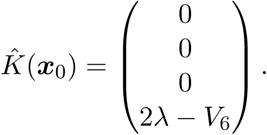

Hence ***x***_0_ is a steady state if we specify

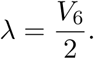

We introduce the Jacobian matrix of 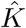 to its variables, evaluated at ***x*** = ***x***_0_,

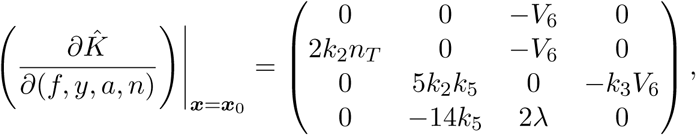

where

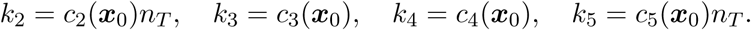

We can easily see that it is invertible. Therefore, by the Implicit Function Theorem, the solution to 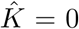 can be locally described as a function of *λ* around 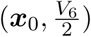.

The curve can be made explicit as a power series expansion in 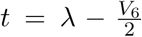, approximating *f, y, a* and *n* around 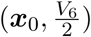. Based on the orders of magnitude in 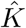 in (24), we expand the variables as

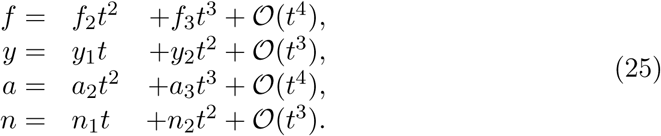

Working out the details (see Supplementary Information; SI) leads us to see the following lowest order coeffcients,

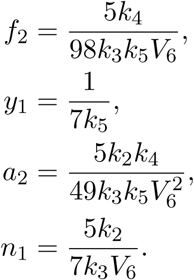

Therefore the curve enters the domain of biologically relevant (***x***, *λ*)-space, for increasing *λ*, because the lowest order terms of the variables are all positive.

In the SI we show that this emergent steady state is locally stable, by checking directly the Routh-Hurwith conditions on the characteristic polynomial for this steady state.

### 3.3 Parameterising steady states by metabolite concentrations

For the following analysis, we will use the matrix *Ñ*_36_ (13), since its Jacobian has most entries have a clear sign.

The steady state equations are given by *Ñ*_36_***v***(***x***) = 0, which can be written out as

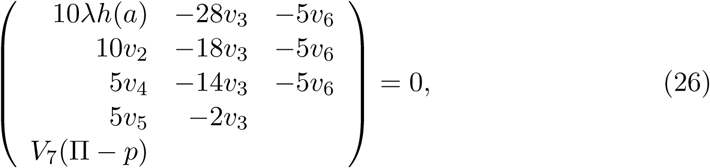

with *v*_2_, *…, v*_6_ as defined in equation (7).

We now focus on regular steady states in which *f, p, y, a, n >* 0 and *a < a*_*T*_, *n < n*_*T*_. The equations

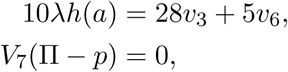

are solved by

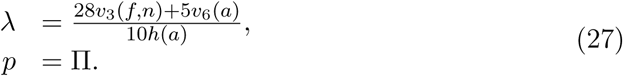

The solution for *p* shows that the pathway neither produces nor consumes phosphate. This is a well-known property and therefore in many other models its concentration is disregarded [19, 7, e.g.]. We include *p* for the imbalanced state, not the steady state, because from experiments we know it is the depletion of inorganic phosphate (*p*) that will restrict the flux of lower glycolysis (*v*_2_), while upper glycolysis (*v*_1_) is unaffected, allowing FBP to accumulate [21].

We aim to find a bifurcation curve in (***x***, *λ*), so the explicit solution for *λ* (27) is already a big step. But its formula is based on a solution ***x***, so equation (27) without further knowledge of ***x*** has no meaning.

The remaining steady state equations (26) are *K*(*f, y, a, n*) = 0, where

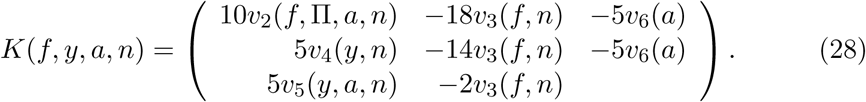

Let *dK* denote the Jacobian matrix of partial derivatives of *K* to *f, y, a* and *n* and note that we can factor it as follows,

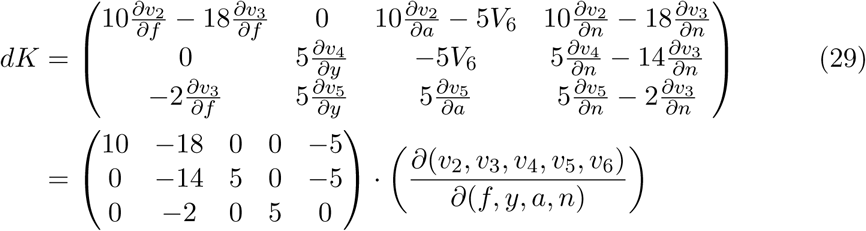

The left matrix is a submatrix of *Ñ*_36_ (12) and we will expand upon the right matrix to reformulate it (30).

With the exception of *v*_1_, the rate functions which make up ***v*** (defined in (7)) are products of individual functions of one variable, in which each function is monotone increasing in its variable (here we include the dependent variables *b* = *a*_*T*_ *- a* and *d* = *n*_*T*_ *- n* for ADP and NAD respectively). Thus any partial derivative of a reaction to a metabolite yields the same product, where the function of the specific metabolite is replaced with its derivative (with a possible minus in front). Moreover, this structure is independent of the specific values of parameters or variables.

We introduce some notation to capture these properties: let 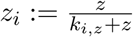. where *z* is a metabolite concentration and *i* follows from which flux *v*_*i*_ we consider. For instance, for flux *v*_3_ we get *v*_3_ = *V*_3_*f*_3_*n*_3_, where 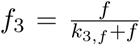 and 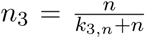. In this notation, 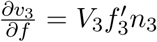, for instance. The possible minus comes from taking a partial derivative to *a* for a function *b*_*i*_ or likewise for *n* and *d*_*i*_, which yields 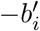 or 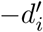 respectively. Then multiplying with 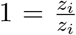 we get that the partial derivative to *z* of a flux is this flux times a logarithmic derivative of its *z*-dependent function,

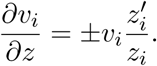

We use here that *z*_*i*_ is nonzero for any concentration *z* and flux *v*_*i*_, which follows from our positivity assumptions on the metabolite concentrations. In this notation, we can rewrite the flux definitions (7) as

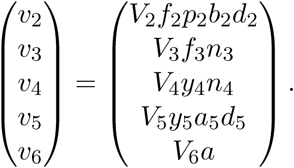

We take partial derivatives to *f, y, a* and *n* and rewrite to get

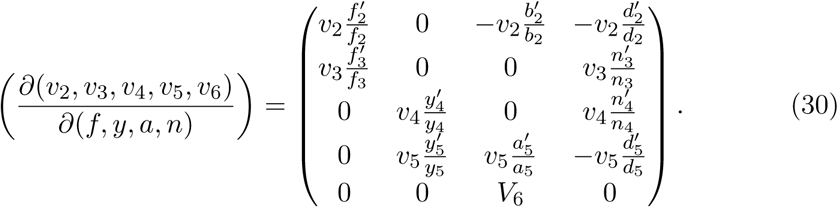

Each of these logarithmic derivatives is positive and the fluxes are also positive. Now we have *dK* in a form amenable to analysis with (29).

The Jacobian *dK* has one column more than it has rows; removing a column and computing the determinant yields a subdeterminant. If a subdeterminant is nonzero for a given solution of *K*(*f, y, a, n*) = 0, then the Implicit Function Theorem (IFT) gives us that the solutions can be locally parameterised in the variable corresponding to the removed column [12]. For example, proving that the subdeterminant where the first column of *dK* is removed is nonzero for a solution (*f, y, a, n*), implies that locally the steady states are on a one-dimensional manifold that can be parameterised by the concentration *f*. We will prove that this is the case.

#### Theorem 5.

*The non-equilibrium steady states can be described as a one-dimensional manifold parameterised by concentration f, given that the following parameter condition holds,*

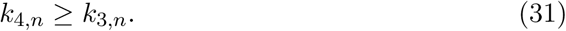

*Proof.* We only need to show that for any solution of equation (28), the subdeterminant of *dK*, where the first column is removed, is nonzero. This implies that the steady states are all locally on a one-dimensional manifold parameterised by *f*, but this extends to a global statement. We can start at an arbitrary solution and follow the locally defined manifold for decreasing *f*. We can continue this until the subdeterminant is zero, but the subdeterminant is nonzero for any non-equilibrium steady state and *f* is decreasing, so we must encounter an equilibrium state, where indeed the subdeterminant is zero. In Section 3.2, we showed that there is only one emergent curve of non-equilibrium steady states from the equilibrium states, hence all steady states are on the same curve, starting at ***x***_0_ for *f* = 0.

So it remains to be shown that the subdeterminant of *dK*, where the first column is removed is nonzero for any solution of equation (28).

Using the reformulation of the Jacobian (30), we can write the relevant subdeterminant as

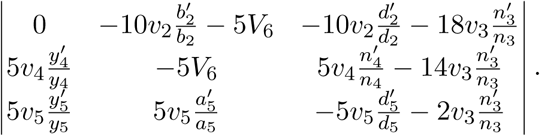

Writing this out, we can see that it is a sum of three negative terms and

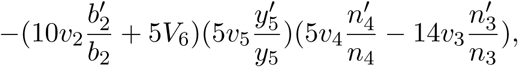

where the right factor does not have a clear sign. We see that the entire determinant is negative if this factor is nonnegative,

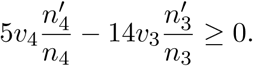

To show the above inequality, we use the steady state equations (26). In particular

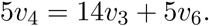

Substituting this equation for 5*v*_4_, we see that the subdeterminant is smaller than zero if the following inequality (or its rewritten form below) holds,

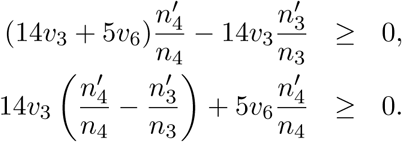

This would follow from the inequality

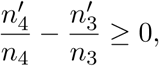

which is a consequence of our assumption *k*_4,*n*_ *≥ k*_3,*n*_ as we can write out the definitions (7),

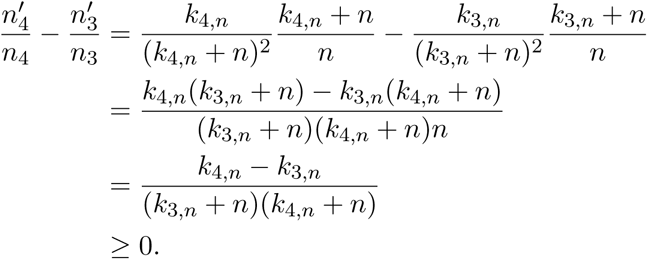

Although the parameter condition of Theorem 5 is comprehensive and acceptable, we can make a more general statement. We now prove that any admissible nonzero vector (*f, y, a, n*) yields a Jacobian *dK* where any two subdeterminants are *never both* zero. In this way we prove that any solution lies on a one-dimensional curve of solutions.

#### Lemma 6.

*Given f, p, y, a, n >* 0 *and a < a*_*T*_, *n < n*_*T*_, *then if a* 3 *×* 3 *subdeterminant of dK is* 0, *then the other subdeterminants are nonzero.*

*Proof.* Assume, for the sake of contradiction, that two arbitrarily chosen subdeterminants are zero. This yields two equations that must hold. We can rewrite these equations as expressions for *V*_2_ and *V*_6_, because in those parameters, the equations are polynomial. Although this is a cumbersome task, it is elementary.

The full computations can be found in the supplementary Mathematica script. The script uses the rewritten form of *dK* from equation (30). The resulting expressions show that *V*_2_ *<* 0 or *V*_6_ *<* 0 follows from the positivity constraints on the other paramters and variables, regardless of their specific values. The parameters *V*_2_ and *V*_6_ are constrained to be positive themselves, so this is a contradiction. We conclude that if one subdeterminant is zero for an admissible solution ***x***, the other subdeterminants are nonzero.

An immediate consequence the Lemma 6 is the following Theorem.

#### Theorem 7.

*All steady state solutions in the interior of the extended metabolite space,* (***x***, *λ*) *>* 0, *lie locally on a smooth one-dimensional manifold.*

### 3.4 The (nonlinear) steady state equations in flux variables

In Section 3.3 we concluded that *f* parameterises all steady state solutions under a mild parameter condition. In this section, we will extend upon this notion of *f* as a flux sensor, by asking whether the steady states can be parameterised by a *flux*, for instance one of the fluxes that consume *f*. Eventually in Theorem 10, we show that flux variable *v*_3_ (the reaction in which FBP is converted into glycerol) can fulfill this role. To obtain this result, we need to solve most equations explicitly. This reduces the five steady state equations to one equations in two flux variables. The Implicit Function Theorem is then applied on this remaining equation to complete the result. We thus first have to eliminate the remaining flux variables, which we manage by setting up a coordinate transform between those other flux variables and the metabolite concentrations.

The hard part is to set up a coordinate transform between three metabolites, *f, n* and *y*, and three reactions, *v*_2_, *v*_3_ and *v*_4_. Matrix *Ñ*36 allows us to construct the steady state equations (26) such that this transformation will reduce the remaining steady state equations *K* = 0 (28) with linear solutions, as *v*_3_ is now a variable of the system and reaction *v*_6_ = *V*_6_*a* is a linear transformation of a variable. Then all that remains to be solved is one equation in two unknowns.

Consider the function,

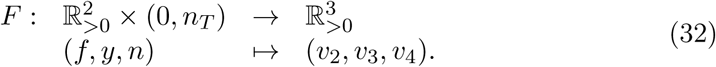

The function *F* is directly based on the reaction rate functions (7); recall their definitions:

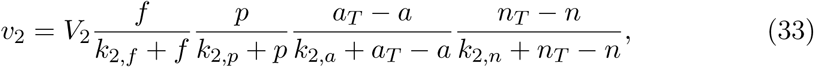

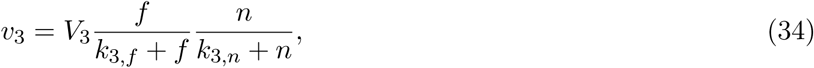

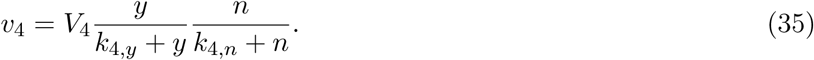

In this section, the variables *a ∈* (0, *a*_*T*_) and *p >* 0 are considered to be parameters of *F*. The variables of *F* model concentrations which, in a living cell, are positive. Therefore, to find steady states in the model that represent a functioning cell at a non-trivial steady state, the domain of *F* is restricted to 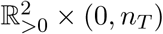.

#### Lemma 8.

*F is smoothly invertible.*

*Proof.* This lemma follows from the Inverse Function Theorem if the determinant of the Jacobian is nonzero for all 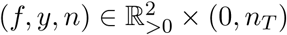.

For any 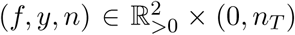, each entry of the Jacobian has a fixed sign. This follows from the monotonicity properties of *v*_2_, *v*_3_, and *v*_4_. The Jacobian

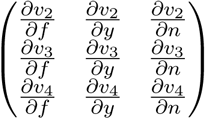

has sign structure

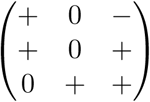

which has strictly negative determinant.

The function *F* is invertible, so it can be used as a coordinate transform. Furthermore we extend *F* with the rescaling *v*_6_ = *V*_6_*a* and the identity on *p* and *λ*, to get a coordinate transform *C* for our phase space by defining

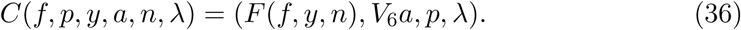

The steady state equations (26) in the new variables are

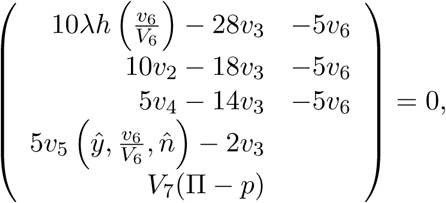

where the inverse coordinate transform *F*^−1^ defines *ŷ* and *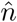* as functions of *v*_2_, *v*_3_ and *v*_4_. Then it can be easily seen that four out of five equations can be explicitly solved,

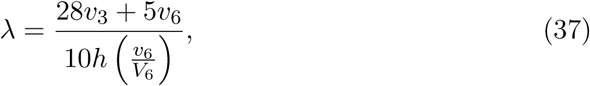

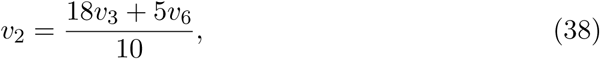

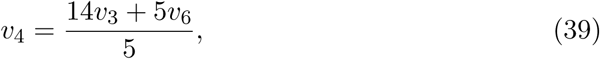

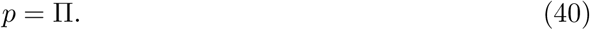

Hence the only steady state equation that is left to solve is

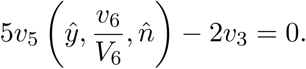

Using the definition of reaction rate *v*_5_ in (7) this reads

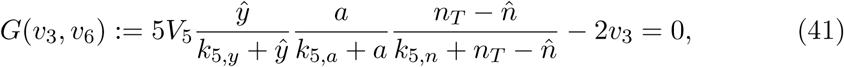

and becomes a nonlinear function of *v*_3_ and *v*_6_ only. To explicitly formulate *G*(*v*_3_, *v*_6_), we need the explicit inverse of *F*, which we will derive below. The exact formulas are found in (49).

Recall the definitions of the rates *v*_2_, *v*_3_ and *v*_4_,

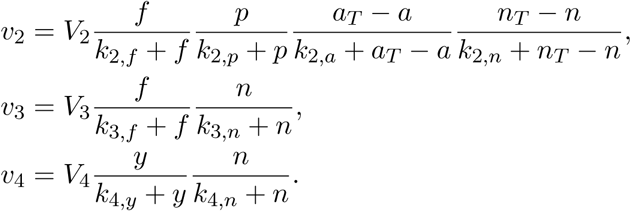

The objective is to define *f, y* and *n* as functions of *v*_2_, *v*_3_ and *v*_4_, for fixed values of *p* and *a*. These functions will be called *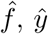*, and *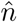*. In fact, we will write *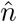* as a function of *v*_2_ and *v*_3_, and then express *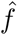* and *ŷ* in *v*_3_, *v*_4_ and *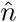*. Let us focus first on this last step, assuming we have computed *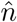* from *v*_2_ and *v*_3_.

The *f* - and *y*-factors in *v*_3_ and *v*_4_ respectively, may be inverted: set

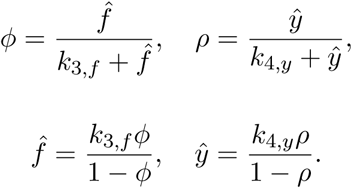

and invert

The expressions for *v*_3_ and *v*_4_ are in this notation given by

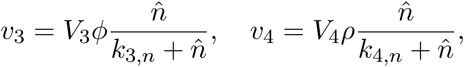

so that

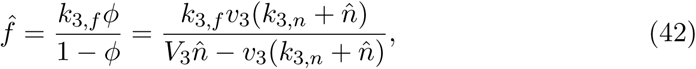

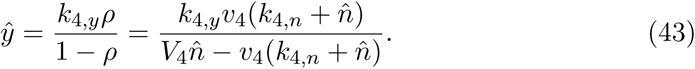

The computation of *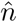* from *v*_2_ and *v*_3_ now follows by considering the definition of *v*_2_. To keep the notation transparent, we introduce

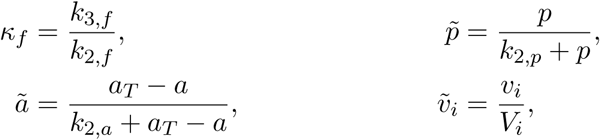

and set

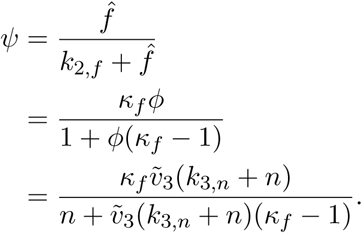

using (42), but with *n* undetermined. Reaction rate *v*_2_ may now be written as

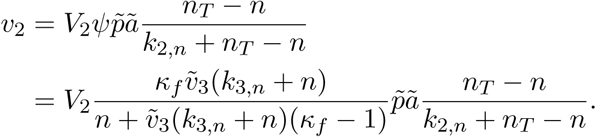

This is a quadratic equation in *n*, written for future reference as

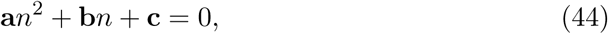

Where

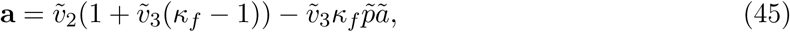

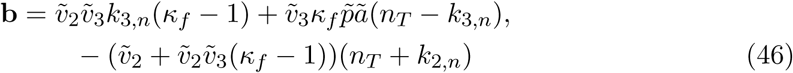

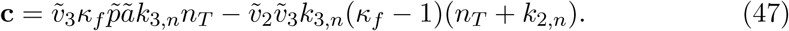

In the SI we show that this equation admits a unique solution *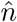* for given values *v*_2_ and *v*_3_ in the range of *F*, namely

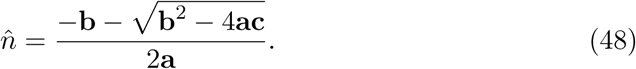

To conclude, the inverse of *F* is explicitly given by

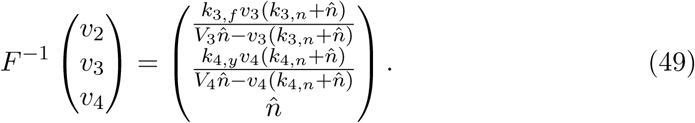

We now use this explicit coordinate transform *F*^−1^ to write the remaining steady state equation *G*(*v*_3_, *v*_6_) = 0. First we define a function *α* of *ρ* and hence of *ŷ* in a similar way as we defined *ψ* as a function of *ϕ* and hence of *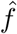*:

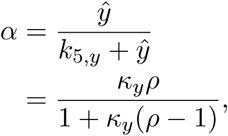

which, using *F*^−1^, becomes

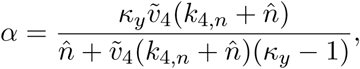

where 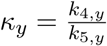. *G* still depends on *v*_4_ (in *α*) and on *v*_2_ (in *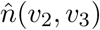*(*v*_2_, *v*_3_)), but using

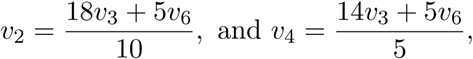

we can now finally write *G* as one function of *v*_3_ and *v*_6_ only,

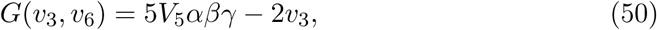

Where

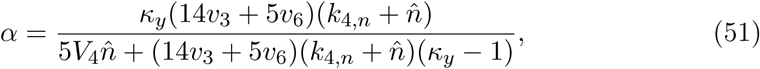

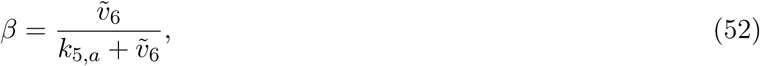

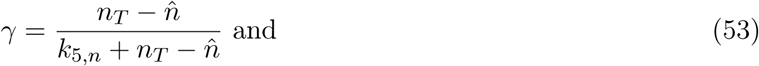

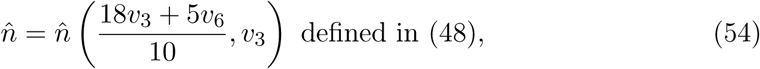

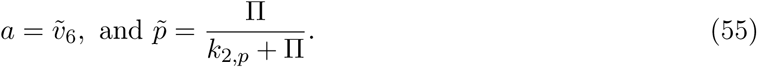

### 3.5 Parameterising the steady states by the glycerol flux

We are now in a position to study *G*(*v*_3_, *v*_6_) = 0, equation (50). With implicit differentiation, we will show that the partial derivative 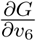 is positive. This result is valid for a broad, but restricted subset of the parameters. Therefore the IFT yields that the solutions to (50) may be expressed as a function *v*_6_ = *H*(*v*_3_). Similar to Theorem 5 we conclude that all steady states are on one curve originating at ***x***_0_ for *v*_3_ = 0 and can be followed until they connect with the boundary of the domain of *F*^−1^. At that point we branch off to the imbalanced states, which we will explore in the next section. (Note that at the boundary of this domain, *v*_3_ is still finite, so the imbalanced states have also become more accessible.)

#### Lemma 9.

*If 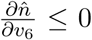 with 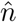 from* (54), *then there exists a function v*_6_ = *H*(*v*_3_), *describing all solutions* (*v*_3_, *H*(*v*_3_)) *to* (50).

*Proof.* The proof uses implicit differentiation. The Implicit Function Theorem yields that if 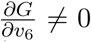 for any (*v*_3_, *v*_6_) in the domain of *G*, then there exists such a function *H*(*v*_3_).

Note that

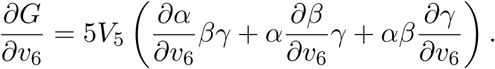

We will show that each of the terms is positive in order to prove 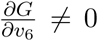 inequalities *α >* 0, *β >* 0, *γ >* 0 follow by definitions (51)–(53), so

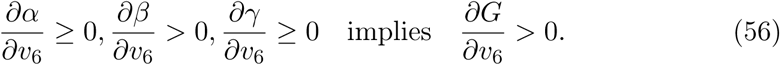

Let

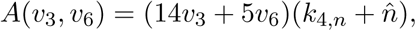

so that

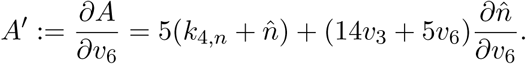

The term *α* as defined in (51) can be rewritten: 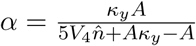

The first term in (56) is shown below.

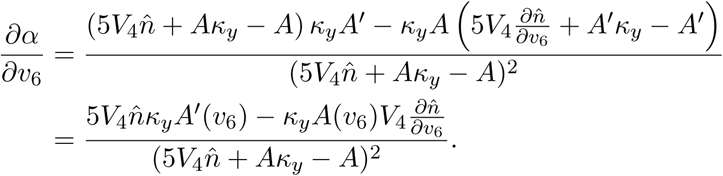

The denominator is positive and we can divide by 5*V*_4_*ff*_*y*_(*k*_4,*n*_ + *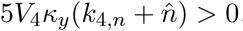*) *>* 0, so the sign of 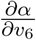 is equal to the sign of

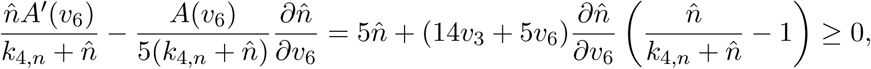

because we have 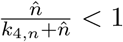 and by assumption 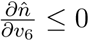.

The second term of (56) is a MM-type function in *v*_6_, so it is monotone increasing.

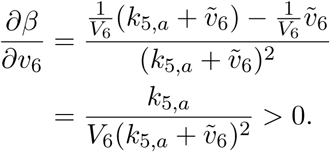

The third inequality follows easily, again under the assumption 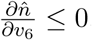,

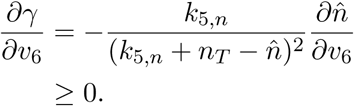

We have now checked all inequalities in (56), which concludes the proof.

#### Theorem 10.

*All solutions to* (10) *in* (***x***, *λ*) *are on a manifold described as a function of v*_3_, *if the following condition on the parameters is satisfied:*

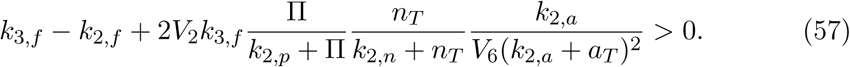

*Proof.* As discussed before, all steady state equations are solved, except (50). Lemma 9 shows that it is suffcient to prove 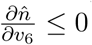 from (54).

For all (*v*_3_, *v*_6_) such that (*v*_2_, *v*_3_, *v*_4_) *∈ D* (*F*^−1^), the defining equation for *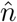* from *F*^−1^ yields 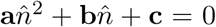. As the left hand side is a function of *v*_3_ and *v*_6_, we know the partial derivative to *v*_6_ is also 0. Hence

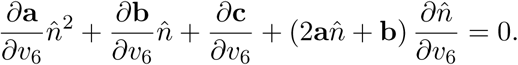

We substitute the solution for 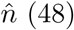 in 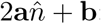:

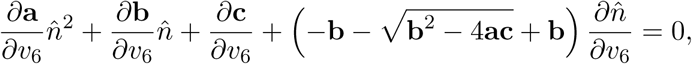

so that

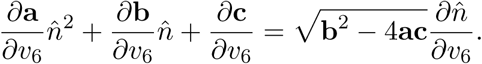

Lemma 8 implies 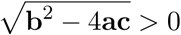, hence the sign of 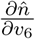 is the same as the sign of

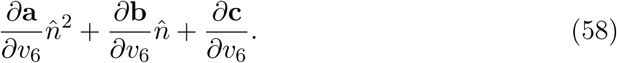

The equations for **a**, **b** and **c**, (45), (46) and (47), written in *v*_3_ and *v*_6_ (and recalling that 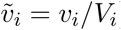), are

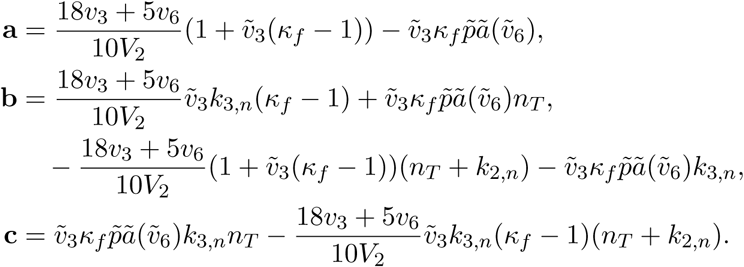

We will show that 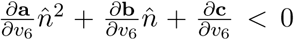. It is suffcient to show it holds for all 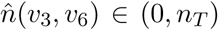. It is first shown that 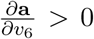 unconditionally. Then as a quadratic function of *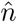*, (58) is negative for all *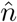* if it is negative at the endpoints, 0 and *n*_*T*_. This restriction at 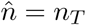 follows unconditionally and the restriction at 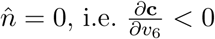, follows precisely from (57), which is why that is a prequisite for this theorem.

The function 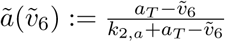 satisfies

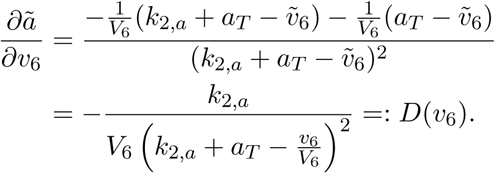

Therefore,

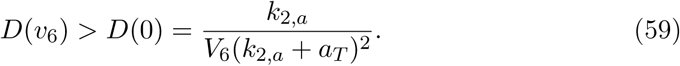

In this notation,

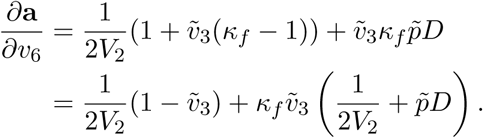

As 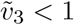 in the domain of *F*^−1^, it follows that 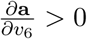.

At 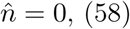 becomes

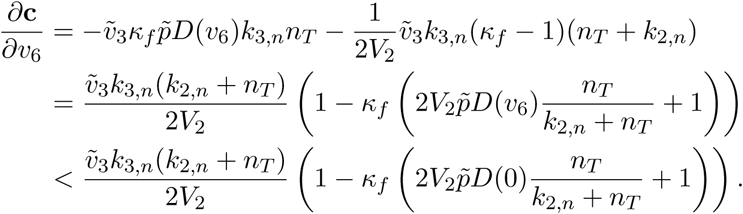

The last inequality follows from the estimate (59) on *D*(*v*_6_). Recall that 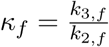 so the sign of 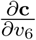 is equal to the sign of

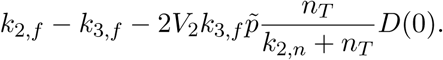

The prequisite for this theorem is (57), which is exactly the above inequality, so it holds that

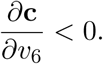

At *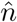* = *n*_*T*_, the formula (58) becomes

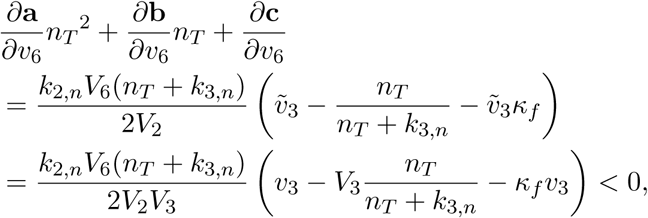

because from (73), 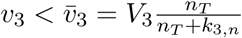.

### 3.6 The imbalanced state

Under certain parameter conditions, glycolysis has been shown to display bistability between a regular steady state, in which the cell functions properly, and an imbalanced state, in which lower glycolysis cannot keep up with upper glycolysis and FBP accumulates [22]. We now turn our attention to this imbalanced state in the core glycolysis model.

The steady states of the model are on a one-dimensional curve (Theorem 7) that at one end always connects to the single equilibrium ***x***_0_ (Section 3.2). Away from the equilibria, the curve continues as the Implicit Function Theorem remains in force. The only possibility for the curve to end is when it connects with the boundary of the restricted domain of *F*^−1^ (32). We will show that in this model, *f* and *y* may both keep increasing to infinity, even simultaneously, depending on the parameter *V*_4_. This extends the analysis on the smaller core model discussed in [14], in which this type of bistability was also studied.

Recall that we introduced in Section 3.4,

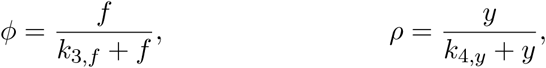

such that

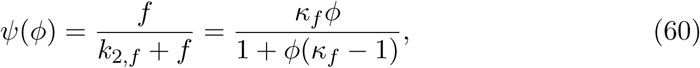

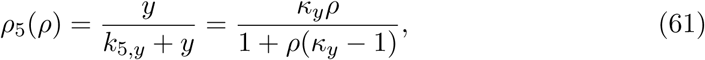

where *ψ*(*ϕ*) and *ρ*_5_(*ρ*) map [0, 1] onto [0, 1], and are monotone increasing, one-to-one functions.

#### Lemma 11.

*If ϕ, ρ ∈* (0, 1] *are fixed, then there exist unique values n*^*\**^ *∈* (0, *n*_*T*_), *a*^*\**^ *∈* (0, *a*_*T*_) *and λ*^*\**^ *>* 0, *and a unique value of V*_4_, *which given by*

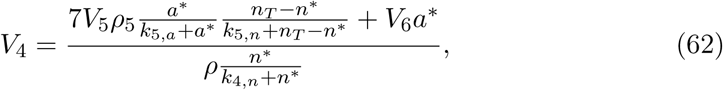

*where* (*ϕ,* π, *ρ, a*^*\**^, *n*^*\**^, *λ*^*\**^) *solves the steady state equations* (10).

*Proof.* This proof is most easily followed from the schematic graphs in Figures 4, 5 and 6.

**Figure 4:**
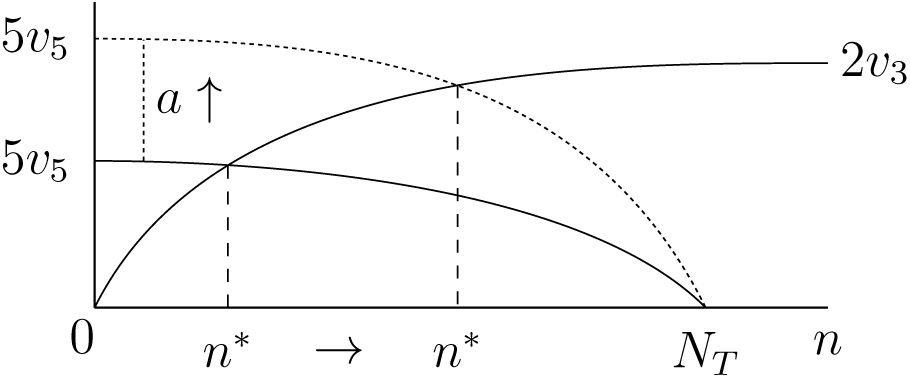
Schematic representation of (64) in terms of *n*. As *a* increases (the dotted graph), *n*^*\**^ is increases.

**Figure 5:**
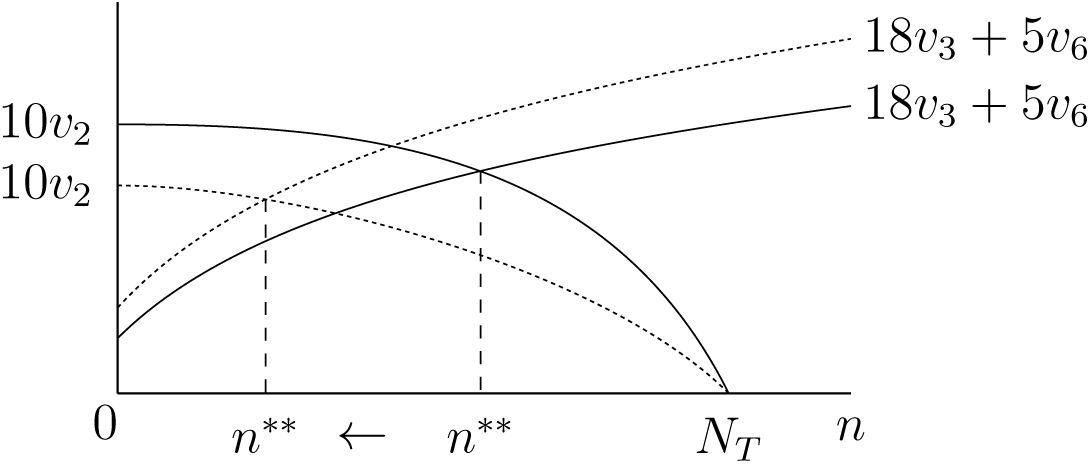
Schematic representation of (65). The dotted graphs are for larger *a*, giving smaller *n*^*\*\**^. Note that *a < a*^*\*\**^ in both cases.

**Figure 6:**
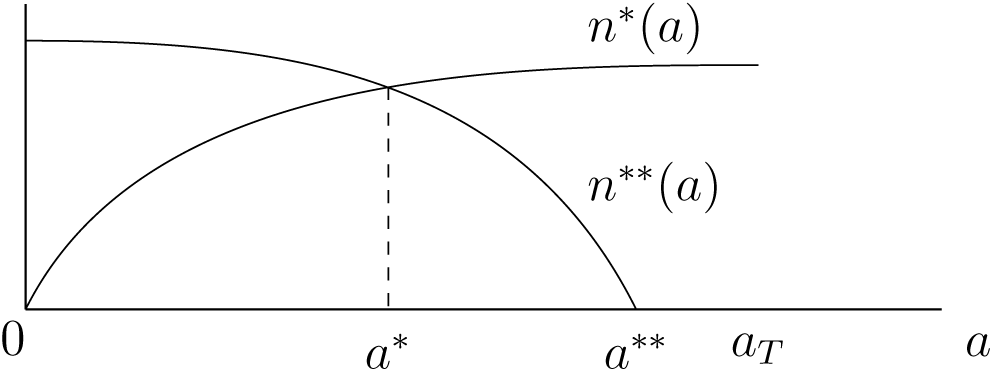
The equations (64) and (65) have a unique solution (*a*^*\**^, *n*^*\**^) as the two solutions for *n* can be solved at the same time with *a* = *a*^*\**^.

Recall the steady state equations can be reformulated using Corollary 2. To prove the lemma, we need all equations to have clear monotonicity properties in the variables *a* and *n*. So depending on what this behaviour in the positive part in an equation is, we either choose *v*_3_ or *v*_5_ together with *v*_6_ to complete that equation. For instance, *v*_2_ is monotone decreasing in *n*, thus in each equation we need the negative parts to be monotone increasing in *n*; since *v*_5_ is monotone decreasing in *n* and *v*_3_ is monotone increasing in *n, v*_3_ is the right choice the argument. This leads us to use the following matrix, where only the third row is different than in the matrix *Ñ*_36_ used in the previous sections,

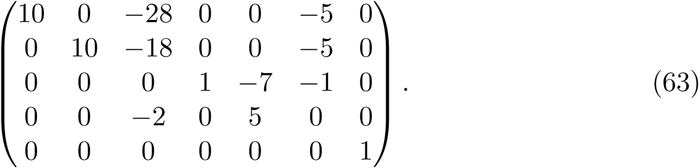

The steady state equations can then be rewritten as

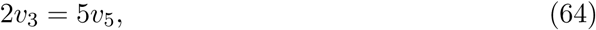

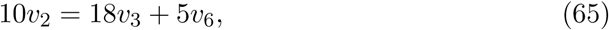

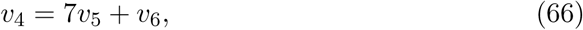

together with

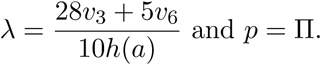

Recall the flux functions (7); written out, the first equation (64) is

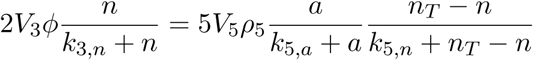

There is a unique solution *n* = *n*^*\**^(*a*): this follows from the IFT, because the lhs is 0 at *n* = 0 and increasing in *n*, the rhs is 0 at *n* = *n*_*T*_ and decreasing in *n*. The function *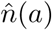* is strictly increasing in *a*, because 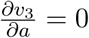 and 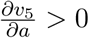(see Figure 4).

The next equation (65) written out is

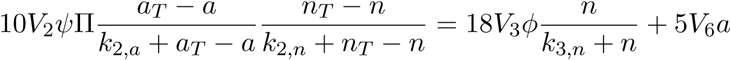

Considered independently from (64), we have a unique solution *n* = *n*^*\*\**^(*a*) if *a* is small enough: note that the lhs is monotone decreasing and the rhs monotone increasing in *n*. At *n* = *n*_*T*_ the lhs is zero and the rhs positive, so we can only use the IFT if, at *n* = 0, there is the inequality

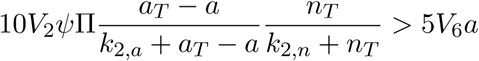

For *a* = 0, this inequality holds and by continuity, it will still hold as *a* increases, with a maximum *a*^*\*\**^, where the inequality becomes an equality, yielding *n*^*\*\**^(*a*^*\*\**^) = 0. Therefore *a ∈* [0, *a*^*\*\**^), yields that *n*^*\*\**^(*a*) *∈*(0, *n*_*T*_) is a solution, which is strictly decreasing in *a*, because 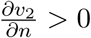, 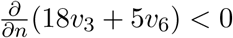 (see Figure 5).

The equations (64) and (65) are not independent, so we need that *n*^*\**^(*a*) = *n*^*\*\**^(*a*). These two solutions of *n* are the same for a unique *a* = *a*^*\**^: the solution to (64) *n*^*\**^(*a*) is 0 at *a* = 0 and increasing, and the solution to (65) *n*^*\*\**^(*a*) is 0 at *a* = *a*^*\*\**^ and decreasing (see also Figure 6).

For simplicity, we will denote *n*^*\**^(*a*^*\**^) as *n*^*\**^. Note that *n*^*\**^ and *a*^*\**^ do not depend on *V*_4_.

The last steady state equation (66) written out with *n* = *n*^*\**^ and *a* = *a*^*\**^ is solved by *V*_4_ *∈* R_+_:

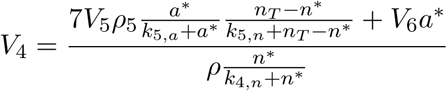

If *V*_4_ is this value, the steady state curve will pass through our given (*ϕ, ρ*) *∈* (0, 1]^2^ with *a* = *a*^*\**^ and *n* = *n*^*\**^ as the remaining variables, and 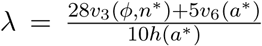, *p* = π.

Note that within this model, it is possible for *f* to accumulate, but also for *y* to accumulate. Accumulation of pyruvate (*y*) is an artifact of this model, and not seen in experiments, since very high pyruvate concentrations impede the production of pyruvate. Accumulation of FBP (*f*), however, does not have a negative effect on its production.

The experimental imbalanced state is characterised as having an accumulation of *f* while the other variables are in steady state. In our analysis, this means that we investigate a steady state of the related model where *ϕ* = 1 is fixed and *f*ff = 0 is *not* part of the steady state equations:

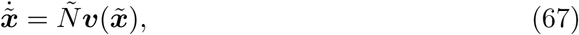

where *ϕ* = 1, ***x***˜ = (*p, y, a, n*) and

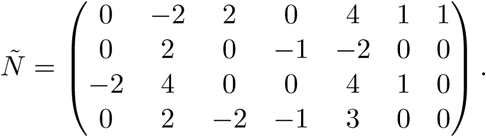

We should however have 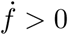 as we want accumulation of *f*.

If we consider the steady states of this problem without assuming *ϕ* = 1, the steady state curve as parameterised by *v*_3_ in Theorem 10 solves these equations, because we have the same equations, without 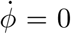. Moreover the solutions were nondegenerate (which follows from Lemma 6), so locally the solutions to *Ñ****v***(***x***) = **0** are a curved plane and the steady state curve separates the part where 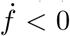 and 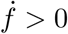. If the steady state curve connects to a point where *ϕ* = 1, this is the starting point of a curve of imbalanced states. We can continue in two directions; one will have 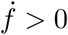 and this is the branch we want to follow.

To find out which branch it is, we manipulate 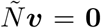. We sum the first and third rows to see that 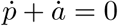, which yields

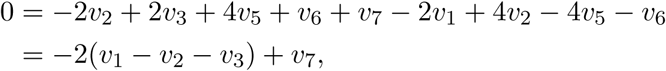

in which we recognise 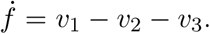. We substitute *v*_7_ = *V*_7_(π *p*) from (7) to get

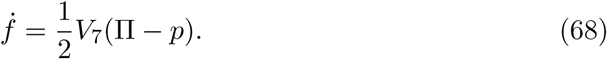

Hence on the curved plane of solutions to 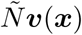, we have that *p <* π is equivalent to 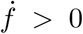, thus we follow the branch for decreasing *p*. This is to be expected biologically: to continue production of FBP, phosphate needs to be added from the vacuole, causing a drop in the vacuole concentration [22].

### 3.7 Numerical illustrations

Parameter-free analysis of the imbalanced state proved too cumbersome. However, Lemma 11 provides us with a good starting point for numerical investigations. We use a custom extension of MATLAB, called MatCont [3] that allows us to continue steady states in a bifurcation parameter. First we simulate our original model (as described in Section 2). Parameter values and initial conditions are given in Table S1.

Lemma 11 yields that for a particular value of *V*_4_ the steady state curve connects to the boundary of the domain of *F*^−1^ at *ϕ* = 1, while *ρ <* 1. In the same way, there exists a critical value *V*_4_ such that we connect at a point where *ϕ <* 1 and *ρ* = 1. Thus manipulation of *V*_4_ should yield both these results. With this in mind, we continue the steady state curve for multiple values of *V*_4_ in MatCont and see that we get a bifurcation: the steady state curve asymptotically goes to *f* = *∞* (*ϕ* = 1), and for increasing *V*_4_ this shifts as it goes asymptotically to *y* =*∞* (*ρ* = 1; see Figure 7).

**Figure 7:**
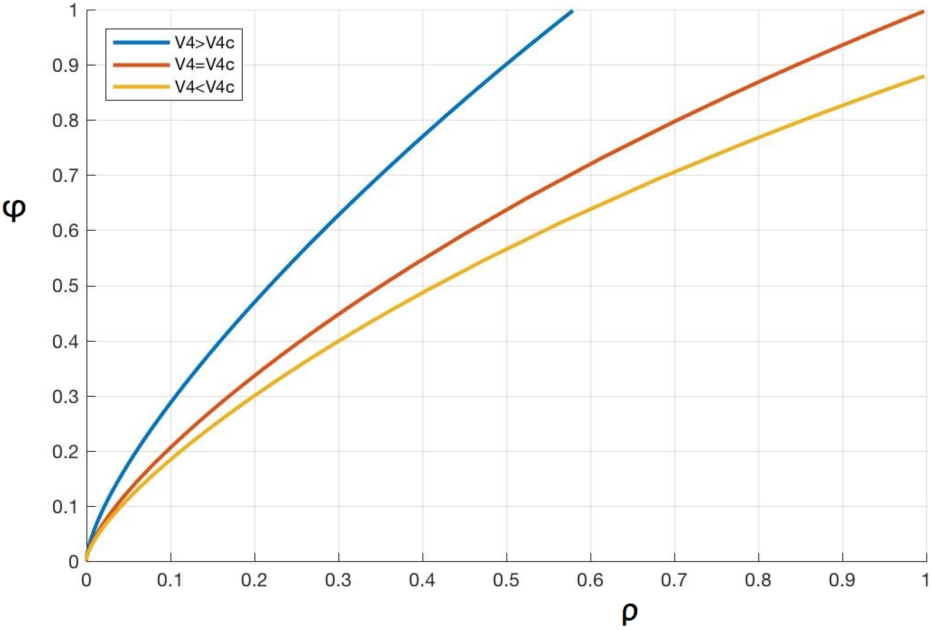
The steady state curve for three different values of *V*_4_. Note that the curve tends to *f → ∞* (*ϕ →* 1) for smaller values of *V*_4_ and to *y → ∞* (*ρ →* 1) for larger values. At the a critical value 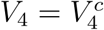, both tend to infinity simultaneously.

To investigate the approach to the imbalanced state, we compactify the dynamics and consider the system with the transformed variable *ϕ* instead of *f*, with time derivative

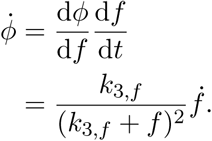

Note 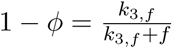, such that 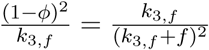 and so

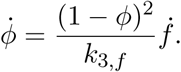

We want to continue the steady states using *λ* as bifurcation parameter to reach *ϕ* = 1. Hence we need to disregard the term 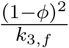 describing the imbalanced states of *ϕ* = 1, and focus on 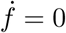.

Using the results from the last simulation, we have a setting in which the steady state curve reaches *ϕ* = 1 while *y* remains finite. Thus we have a starting point ***x***_*∞*_ from which we can continue and follow the imbalanced state. This is found by solving the system “at infinity” (67) for decreasing *p*: since 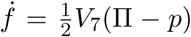(68), *p <* π means 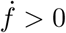. Parameter *V*_7_ in *v*_7_ does not influence the regular steady states, but may be varied in the imbalanced states to produce bistability [14].

The combined results of the last two sections yield a steady state curve and a curve of imbalanced states (see Figure 8). The steady state curve has increasing *λ* and has negative eigenvalues, making it locally stable. The imbalanced state continues in increasing *λ* and is also stable, until there is a limit point near *p* = 0, where we get rapidly decreasing *λ* and instability, followed by another limit point to increasing *λ* even closer to *p* = 0 (see Figure 8). Thus with *λ* a bit greater than its value in this second limit point the system has a stable steady state and a stable imbalanced state. This is an example of the bistability between a regular and an imbalanced state as found experimentally.

**Figure 8:**
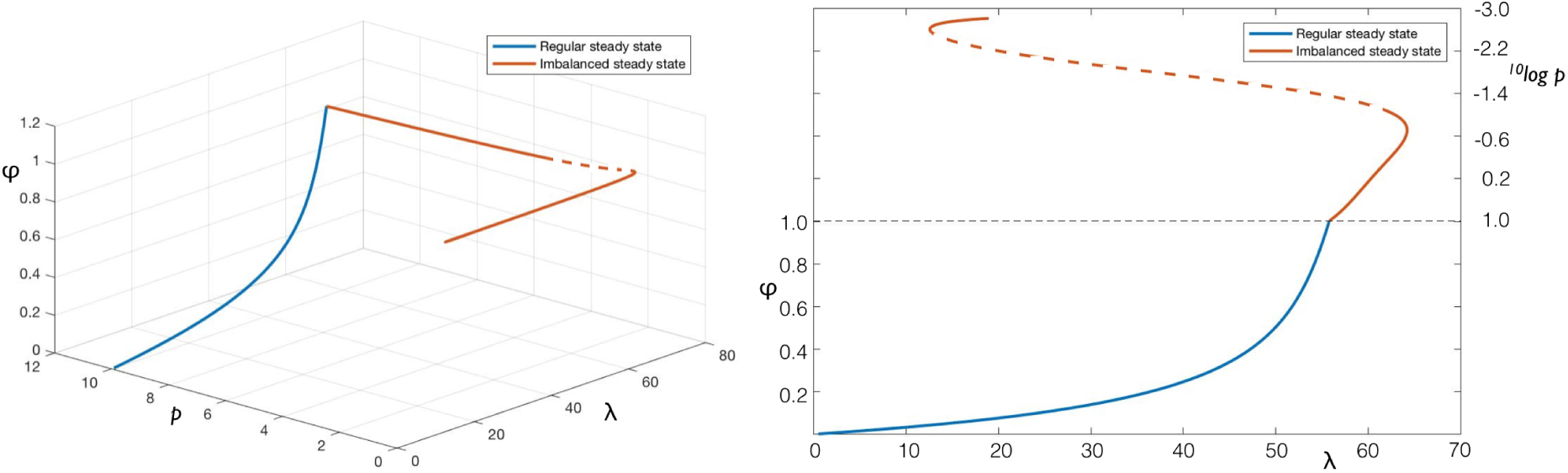
The steady state curve for the transformed variable *ϕ,* reaching *ϕ* = 1 (in blue) and the imbalanced states branching off from this (red). Left: bifurcation curves in (*p, λ, ϕ*)-space; Right: the same curves, showing the *p* and *ϕ* values along the curve, to better illustrate the bistability between regular and imbalanced states.

## 4 Discussion

With the goal of parameter-free analysis for more complex metabolic pathways, we have given a detailed description of the steady states of the core model of yeast glycolysis described in Section 2. It was possible to prove under very mild parameter conditions that FBP parameterises the steady states, but to show a similar statement for the glycerol flux variable *v*_3_ turned out to be much more involved. Also the analysis of imbalanced states proved partially beyond current techniques.

### 4.1 A surprisingly simple condition for non-equilibrium steady states

If the parameter condition *V*_1_ *> V*_6_*/*2 is met, the equilibrium state is unstable and the emerging steady state is stable (Section 3.2). In other words, if the rate of consumption of ATP (*V*_6_) is less than twice the rate of glucose feeding into glycolysis, then the cell can balance the production and consumption of ATP in a stable steady state. This ratio of 1 : 2 is exactly the ratio between the reactions *v*_1_ and *v*_6_ in the Elementary Flux Mode *EFM*_1_, which represents the normal glycolytic flux through the pathway (11). This does not even take into account the other EFM, *EFM*_2_, which does involve the side branches and should account for a part of the glucose uptake, specifically the part that does not produce any ATP (as seen by *v*_6_ = 0 in this flux vector). This is surprising: as soon as the rate of glucose uptake can support the ATP consumption downstream, the system has an emerging stable steady state with balanced metabolism. So although the condition *V*_1_ *> V*_6_*/*2 would seem to be a bare minimum for stability, it is all that is required.

The subtle point in our proof where we see why this bare minimum is enough is in the power series expansion (25). If we insert this expansion in the reactions (7), we get exactly that the nonzero reactions in *EFM*_1_ are of order *t*^2^, while the side branch reactions *v*_3_ and *v*_5_ are of order *t*^3^. Hence, near the equilibrium, the flux directly through glycolysis is dominating the glucose intake.

This result is also simpler in the more complicated model studied in this paper than in the previous simpler core model studied in [14]. In that smaller model, there was a richer set of solutions. The addition of NADH/NAD householding, which was lacking from [14], induced a more complicated model with two instead of one side branch (the other being the pyruvate to succinate branch, which was specifically added for redox balance) and simplified the bifurcation structure considerably.

### 4.2 The one-dimensional curve of non-equilibrium steady states

The steady states are locally a one-dimensional manifold, by Theorem 7. So it follows that the only ways the steady state curve could change character is if a pair of eigenvalues passes the imaginary axis (Hopf bifurcation) or if the curve changes direction in the parameter *λ* (fold bifurcation). We have neither found nor excluded these types of bifurcations in our model, but the main result of our work is that these are the only possibilities. This means that there are no pitchfork bifurcations, which would create multiple steady state branches, and no other transcritical bifurcations. In case of a fold, the steady state would become unstable, but it would coexist with a stable steady state as the curve changes direction. In case of a Hopf bifurcation, a limit cycle would emerge, possibly leading to oscillations, a well-known phenomena in yeast glycolysis [5], and also present in the previous core model we extended here [14]. If in addition it would be possible to show that the curve is monotone increasing in the parameter *λ*, then all the points on the curve have different *λ*, so for a fixed *λ*, the steady state is unique. We have not been able to prove this monotonicity property of the curve, but we found it in all the numerical simulations.

### 4.3 FBP as a flux sensor

As introduced at the start of our work, we wanted to understand if FBP always parameterises the steady state curve, so that it could function as a flux sensor [2, 8]. Indeed, in Theorem 5 this was shown to be true if

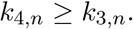

Note that this is a sufficient condition and could still be true if *k*_4,*n*_ *< k*_3,*n*_. The inequality above is in correspondence with measured values of the affinities of NADH to the two reactions in which FBP is a substrate [20]. This suggests that in all likelihood FBP could indeed act as a flux sensor under practically any parameter setting; in other words, this fact is explained by the stoichiometry and structure of the kinetics of the rate functions of this metabolic pathway, rather than by the precise parameters involved in those rate functions.

Another question was whether a flux could parameterise the steady states, which we found can be shown to be true for the glycerol flux *v*_3_ in Theorem 10. However, we needed a restriction on the parameters (57). This restriction involves many parameters, but most notably *k*_2,*f*_ and *k*_3,*f*_. A suffcient condition for (57) to hold is that *k*_2,*f*_ *< k*_3,*f*_, which should be interpreted as that lower glycolysis has a higher affinity for FBP than glycerol production. This is a very reasonable assumption, and is in line with experimental data [20]. The central flux through glycolysis is the major flux in our model, therefore its response to a change in FBP should be more sensitive than the response of a side branch.

### 4.4 The imbalanced state

Experimentally, the imbalanced state occurs in a fraction of the cells [22], and there is a bistability between steady and imbalanced states; the dynamics converge to either stable state depending on the initial conditions. Most initial conditions will approach a stable steady state, but there is a small basin of attraction for the imbalanced state. This matches well with the numerical results as depicted in Section 3.6.

The continuation of the imbalanced state (the red curve in Figure 8) consists of three parts, due to two fold bifurcations. We see that for a certain range of the parameter *λ*, the stable steady state and two imbalanced states coexist, where one imbalanced state is stable and one is unstable. For a fixed *λ* in this range, the stable imbalanced state is very close to zero for the concentrations *a* and *p*, which coincides with the experimental picture: the cell depletes the inorganic phosphate (*p*) and ATP (*a*) concentrations in the cell, and all the production lower glycolysis can muster is immediately invested to produce more FBP, which therefore accumulates in the cell [22].

### 4.5 Scope of the techniques and general outlook

In this paper we have used three main techniques to study the bifurcation structure of the moderately detailed pathway with explicit kinetics: EFMs, the Implicit Function Theorem, and coordinate transformations. Here we briefly discuss the generality of our approach to other (and larger) pathways.

The use of EFMs to construct alternative steady state equations (Proposition is general. This method will be of use particularly for models with between four to about ten independent variables; with less, one can oversee the recombination of rows easily, and with more equations there will be a combinatorial explosion in the number of EFMs, and they do not provide additional insight. For higher numbers of independent variables, Extremal Pathways might be more suitable than EFMs, as there are less of those, but at some point also these will become cumbersome to use.

The Implicit Function Theorem was used especially to prove that the nonequilibrium steady states formed a single curve emerging from one equilibrium steady state. The technique uses smaller subdeterminants of the complete Jacobian matrix, and the linearity of *V*_max_ parameters in reaction functions to prove that at least one subdeterminant is always nonzero. This technique scales in principle to much larger networks, and it should eventually be possible to prove in full generality whether a detailed model such as the ones by Teusink *et al.* [20] or Hynne *et al.* [9] have the same property.

Finally, we used coordinate transformations to reformulate the steady state equations, for two reasons: to exploit the linearity of fluxes in those equations, and to compactify the dynamics and study imbalanced states “at infinity”. In our case the inverse transformation could be explicitly calculated, but this is not to be expected for larger networks (unless they involve several smaller individual transformations between metabolites and fluxes). Moreover, the amount of work necessary to prove that *v*_3_ parameterises the steady states was considerable, and was strongly dependent on the explicit choices of reaction rate functions, their product structure and whether they happened to be increasing or decreasing in their respective variables. We do not expect that such calculations are possible in much larger networks. On the other hand, setting up the transformation itself was straightforward, and already allowed the study of both regular steady states and imbalanced states.

## Acknowledgements

GO and RP gratefully acknowledge funding on grant NDNS+ 613.009.012. GO would also like to thank Sander Hille for his guidance during early ventures into this research.

## Supplementary Information

The supplementary Mathematica file contains the calculations for

- The expansion in equation (25), Section 3.2.
- The local stability of steady states emerging from the equilibrium ***x***_0_ at the end of Section 3.2;
- The calculations in the proof of Lemma 6, dSection 3.3.

In Section 3.4, we mentioned that equation (44) admits a unique solution. The proof is given here.

We need to show that *n*^*\**^ is uniquely determined by *v*_2_ and *v*_3_ in the image of *F*. Recall the quadratic equation *n*^*\**^ needs to solve,

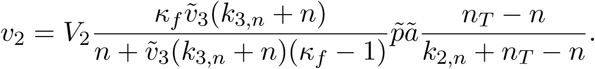

A quadratic equation can have zero, one or two solutions, but since *F* is smoothly invertible on its domain (Lemma 8) we know that there has to be exactly one on the restricted domain of *F*.

We recall the short version of this equation,

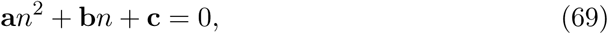

Where

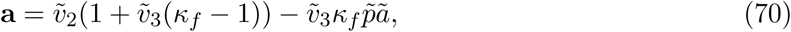

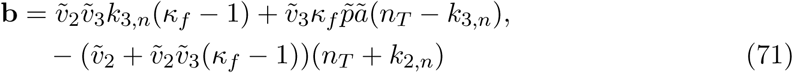

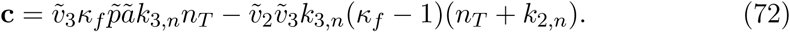

The transformation *F* is bounded. The borders of the image in (*v*_2_, *v*_3_, *v*_4_)-space are complicated curves, but each reaction has a clear global upper bound and 0 as a lower bound: the typical Michaelis Menten fraction 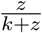 approaches 1 for *z → ∞*. Furthermore *n ∈* (0, *n*_*T*_), so we have the following bounds for *F*,

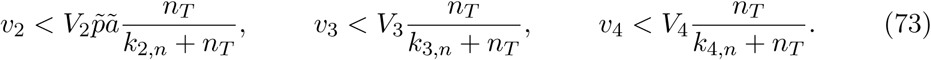

Denote these upper bounds for *v*_*i*_ as 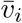.

The transformation *F* has this bounded image, which yields a bounded domain for *F*^−1^.

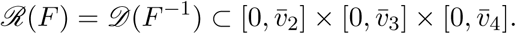

This will be a tool to prove which of the two roots of the quadratic equation for *n* is the right one.

### Lemma 12.

*For all v*_2_ *and v*_3_, ***c*** *from* (72) *is positive.*

*Proof.* From (73), we have 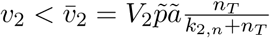 Now (72) can be rewritten to a form that is positive:

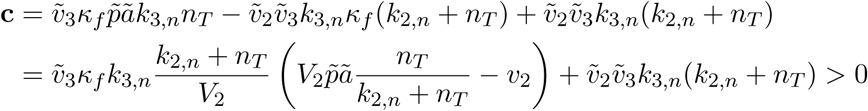

### Lemma 13.

*If* ***a*** *from* (70) *is positive, then 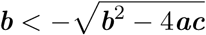.*

*Proof.* This implication follows from a sign argument. The term **c** is positive by Lemma 12 and **a** *>* 0 is assumed. Therefore **b**^2^ - 4**ac** *<* **b**^2^. Note *F* is one-to-one, implicitly shown in Lemma 8, so there is a single solution to (69) that is in the interval (0, *n*_*T*_). It follows that the discriminant **b**^2^ - 4**ac** is positive, because its square root has to be real. Now 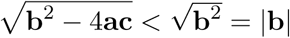. A positive **b** is excluded as both roots of (69) are then negative, hence 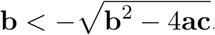.

### Lemma 14.

*The relevant solution to* (69) *is*

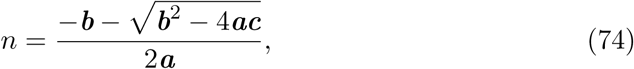

*with continuous extention* 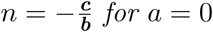.

*Proof.* There are three cases.

**a** *>* 0. By Lemma 13, 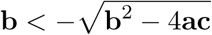. So both solutions are positive,

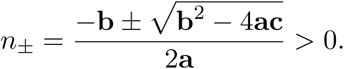

By Lemma 8 there is only one solution in*v*(0, *n*_*T*_), so *n*_*T*_ must separate the two solutions. Of these, *n*_-_ is closer to 0 as 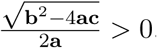.

**a** *<* 0. In this case, we have *n*_+_ *<* 0. From Lemma 12, **c** *>* 0, so 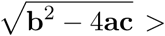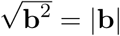, therefore

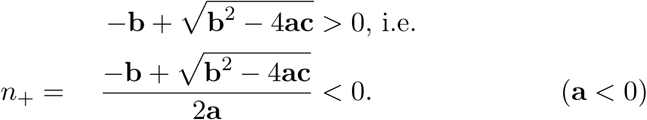

**a** = 0. Now (69) becomes 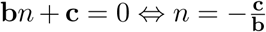. This is a continuous extention of *n*_-_:

**c** *>* 0 by Lemma 12. If **a** is near 0, then (69) is close to **b***n* + **c** = 0, so **b** *≥* 0 would imply that *n >* 0 can only be a solution to (69) if **a** *<* 0 and *n* is large, then as **a** approaches zero, this value would have to pass *n*_*T*_ making all solutions irrelevant. Hence we have **b** *<* 0.

From this it follows that 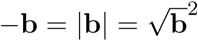. Now we can show that indeed *n* may be continuously extended:

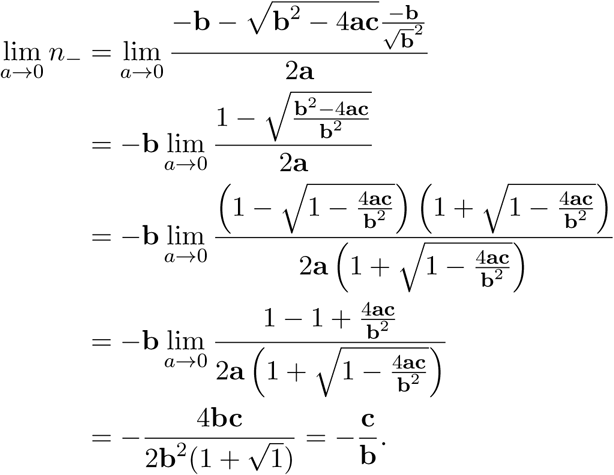

So *n*_-_ is the relevant solution.

**Table 1:**
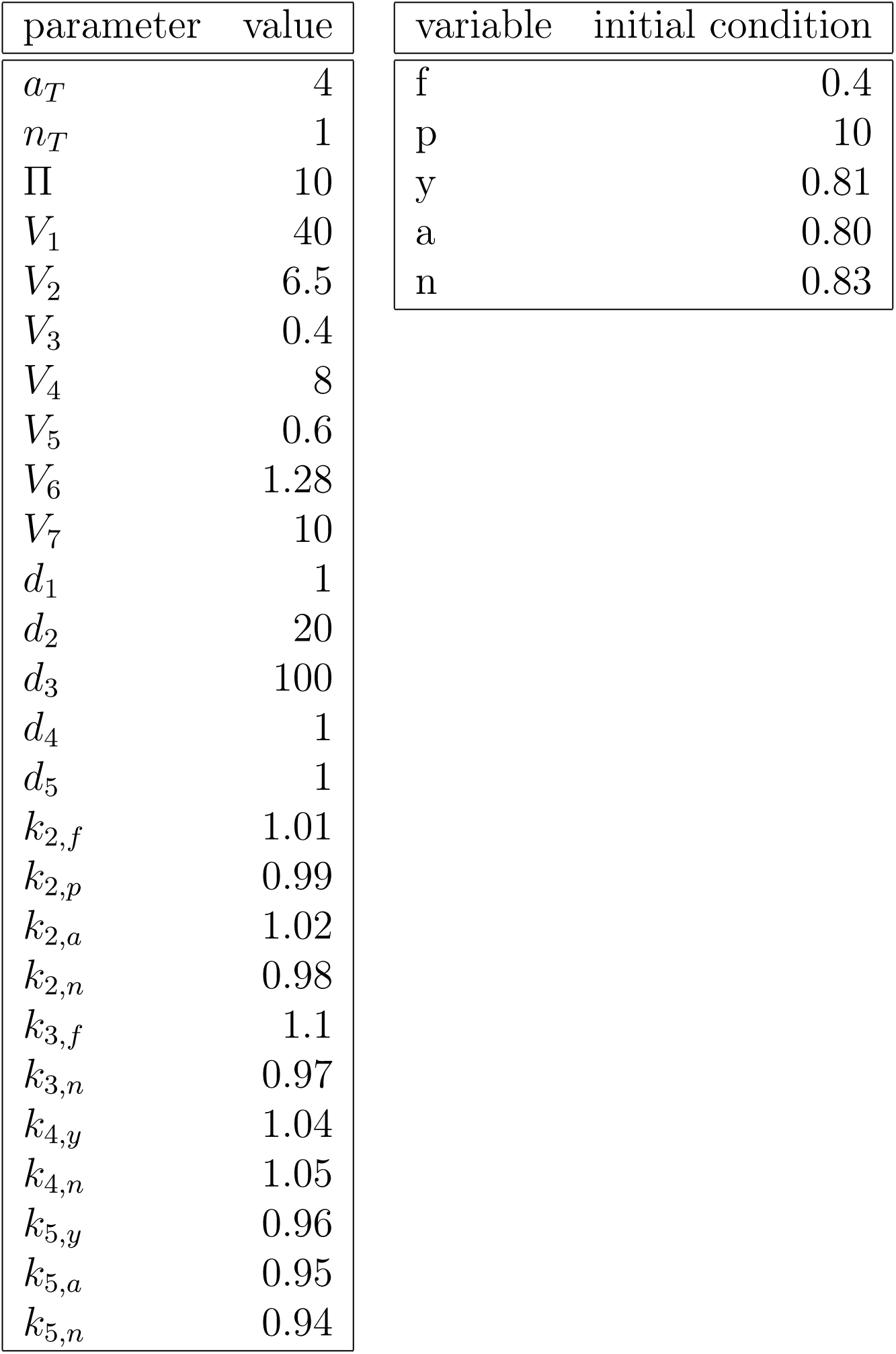
Parameter values and initial conditions used in MatCont simulations.

